# Comprehensive characterization of multi-omic landscapes between gut-microbiota metabolites and the G-protein-coupled receptors in Alzheimer’s disease

**DOI:** 10.1101/2022.09.20.508759

**Authors:** Yunguang Qiu, Yuan Hou, Yadi Zhou, Jielin Xu, Marina Bykova, James B. Leverenz, Andrew A. Pieper, Ruth Nussinov, Jessica Z.K. Caldwell, J. Mark Brown, Feixiong Cheng

**Affiliations:** Genomic Medicine Institute, Lerner Research Institute, Cleveland Clinic, Cleveland, OH 44195, USA; Department of Molecular Medicine, Cleveland Clinic Lerner College of Medicine, Case Western Reserve University, Cleveland, OH 44195, USA; Lou Ruvo Center for Brain Health, Neurological Institute, Cleveland Clinic, Cleveland, OH 44195, USA; Harrington Discovery Institute, University Hospitals Cleveland Medical Center, Cleveland, OH 44106, USA; Department of Psychiatry, Case Western Reserve University, Cleveland, OH 44106, USA; Geriatric Psychiatry, GRECC, Louis Stokes Cleveland VA Medical Center; Cleveland, OH 44106, USA; Institute for Transformative Molecular Medicine, School of Medicine, Case Western Reserve University, Cleveland 44106, OH, USA; Department of Neuroscience, Case Western Reserve University, School of Medicine, Cleveland, OH 44106, USA; Computational Structural Biology Section, Frederick National Laboratory for Cancer Research in the Cancer Innovation Laboratory, National Cancer Institute, Frederick, MD 21702, USA; Department of Human Molecular Genetics and Biochemistry, Sackler School of Medicine, Tel Aviv University, Tel Aviv 69978, Israel; Lou Ruvo Center for Brain Health, Neurological Institute, Cleveland Clinic, Las Vegas, NV 89106, USA; Department of Cardiovascular and Metabolic Sciences, Lerner Research Institute Cleveland Clinic, Cleveland, OH 44195, USA; Center for Microbiome and Human Health, Lerner Research Institute, Cleveland Clinic, Cleveland, OH 44195, USA; Case Comprehensive Cancer Center, Case Western Reserve University School of Medicine, Cleveland, OH 44106, USA

## Abstract

Accumulating evidence suggests that gut-microbiota metabolites contribute to human disease pathophysiology, yet the host receptors that sense these metabolites are largely unknown. Here, we developed a systems pharmacogenomics framework that integrates machine learning (ML), AlphaFold2-derived structural pharmacology, and multi-omics to identify disease-relevant metabolites derived from gut-microbiota with non-olfactory G-protein-coupled receptors (GPCRome). Specifically, we evaluated 1.68 million metabolite-protein pairs connecting 408 human GPCRs and 516 gut metabolites using an Extra Trees algorithm-improved structural pharmacology strategy. Using genetics-derived Mendelian randomization and multi-omics (including transcriptomic and proteomic) analyses, we identified likely causal GPCR targets (C3AR, FPR1, GALR1 and TAS2R60) in Alzheimer’s disease (AD). Using three-dimensional structural fingerprint analysis of the metabolite-GPCR complexome, we identified over 60% of the allosteric pockets of orphan GPCR models for gut metabolites in the GPCRome, including AD-related orphan GPCRs (GPR27, GPR34, and GPR84). We additionally identified the potential targets (e.g., C3AR) of two AD-related metabolites (3-hydroxybutyric acid and Indole-3-pyruvic acid) and four metabolites from AD-related bacterium *Eubacterium rectale*, and also showed that tridecylic acid is a candidate ligand for orphan GPR84 in AD. In summary, this study presents a systems pharmacogenomics approach that serves to uncover the GPCR molecular targets of gut microbiota in AD and likely many other human diseases if broadly applied.

## Introduction

Recent advances in chemogenomic and pharmacogenomic approaches have shown that G-protein-coupled receptors (GPCRs) mediate a significant portion of host-microbiota interactions^1, 2^. Indeed, broad distribution of GPCRs in the gastrointestinal tract and brain may underlie their importance in shaping the “gut-brain axis” and other aspects of host pathophysiology^3, 4^. Of those, over 100 receptors are “pharmacologically dark” or “orphan” GPCRs, whose endogenous ligands are unknown and their physiological roles have not been well-investigated^5^. Several gut metabolites have been recognized as agonists of orphan GPCRs from functional screening^1, 6–9^. For example, orphan GPR35 is activated by 5-hydroxyindoleacetic acid in neutrophil recruitment^10^, and orphan GPR84 is agonized by medium-chain fatty acids in inflammation^11^. However, accurate structures of GPCRs, especially for orphan GPCRs, are lacking. This limits the power to identify metabolite-GPCR regulation. Recent advances in computational sciences and structural biology, such as Alphafold2^12^, offer unprecedented opportunities to investigate the relationship between structure and function, in particular for orphan GPCRs without known structures.

Alzheimer’s disease (AD) is a progressive neurodegenerative disorder caused by multiple pathophysiological factors, including both genetic and environmental factors^13, 14^. Without long-term outcomes for clinical prevention strategies and disease-modifying treatments that slow the neurodegenerative process, recent estimates predict that there will be more than 13.8 million people with AD living in the United States by 2050^13^. The environmental exposome, which is the measure of all the exposures of an individual across their lifetime, plays crucial roles in the etiology and progression of AD^15, 16^. Withing this domain, alterations in the gut microbiome and microbe-associated metabolites is increasingly regarded as an important non-heritable neural exposome associated with AD pathogenesis and progression^17–19^.

Growing evidence shows that patients with AD dementia or mild cognitive impairment have microbial dysbiosis characterized by enterotype diversity and abundance^20, 21^. Recent studies have also suggested that alterations of microbiota could contribute to AD-related pathologies in both central and peripheral neural systems^15, 22^. For example, microbiota depletion attenuates inflammation and brain pathology in the APP/PS1 transgenic mouse model of AD^23^, and aging-associated neurocognitive and immune decline can be ameliorated by a fecal microbiota transplant from young to aged mice^24^. Advanced metabolomic approaches have recently identified that both trimethylamine N-oxide^25^, δ-valerobetaine^26^ and N^6^-carboxymethyllysine^27^ are elevated in AD pathobiology. However, the underlying molecular mechanisms linking gut metabolites to human disease, including AD, remain largely unknown. This is a critical area of unmet need, as the identification of “gut-brain axis”^28^ interactions holds potential for fostering discovery and development of new therapeutic approaches for AD. In particular we are only beginning to understand how gut microbe-derived metabolites engage host receptor systems to shape human health and disease.

In this study, we present a systems pharmacogenomics framework that uniquely integrates machine learning and network-based genetics approaches and multi-omics findings to prioritize potential therapeutic targets from GPCRome and microbiota-derived metabolite-GPCR relationships. We systematically evaluated over 1.68 million metabolite-GPCR pairs using AlphaFold2-derived computational biophysical approaches and machine learning models, which enabled genetics-derived Mendelian randomization and multi-omics (including transcriptomic and proteomic) analyses leading to the identification of potential druggable GPCR targets (including orphan GPCRs) for AD. This pharmacogenomic framework thus offers a powerful tool to identify gut metabolite-based therapies via targeting human GPCRs for treating AD. In principle, this approach could be applied productively to other human diseases if broadly applied.

## Results

### A systems pharmacogenomics framework identifies associations between gut-microbiota metabolites and the GPCRome

We constructed a systems pharmacogenomics framework to identify associations between microbiota-derived metabolites and the GPCRome using AD as a prototypical example. This framework entails four steps (Fig. 1a): (1) Identification of likely causal GPCRs in AD using Mendelian randomization analysis; (2) Prioritization of AD-associated GPCRs from multi-omics (including AD brain transcriptomics and proteomics) analysis; (3) Characterization of microbial metabolites from human gut strains; and (4) Identification of associations between metabolites and the GPCRome using AlphaFold2-derived computational biophysical approaches and an Extra Trees model. Specifically, we re-constructed structural models of 408 non-olfactory GPCRs by using AlphaFold2, including 124 orphan GPCRs (Fig. 1b). We next developed an Extra Trees model by leveraging three-dimensional (3D) interaction features derived from large-scale metabolite-GPCR complexes from molecular docking experiments, followed by reconstruction of the metabolite-GPCR interactome network using the Extra Trees model-predicted binding affinity. Finally, we investigated the mechanism-of-actions of several AD-relevant gut metabolites that potentially treat AD via specifically targeting disease-associated GPCRs.

**Fig. 1.**
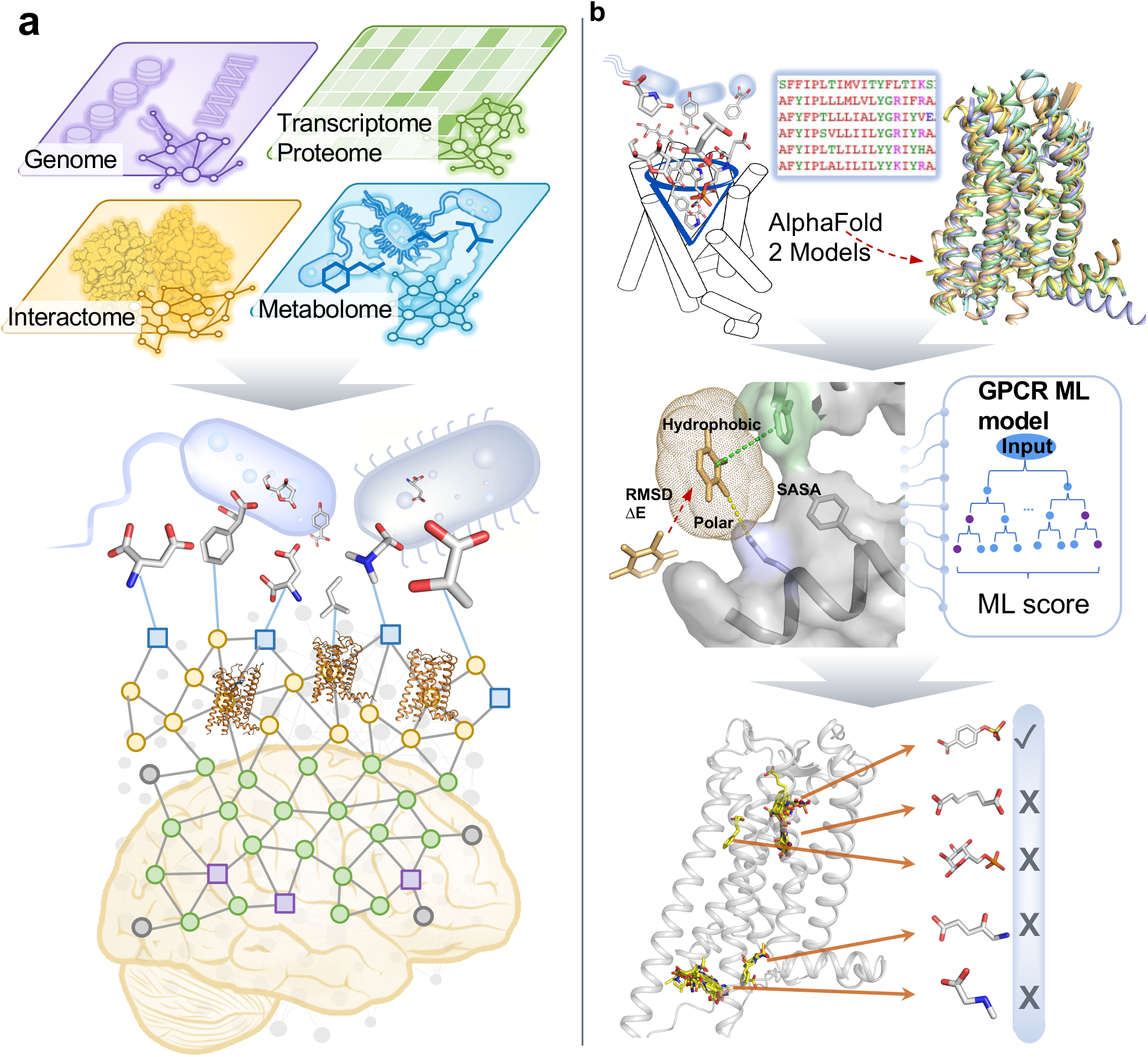
Schematic of a systems pharmacogenomics framework illustrating associations between gut-microbiota metabolites and GPCRome. **a** Schematic diagram illustrating associations between gut-microbiota metabolites and GPCRome in AD. Specifically, AD related genes, proteins, microbiota-derived metabolites datasets were compiled from multi-omics data including genome (purple), transcriptome/proteome (green), metabolome (cyan) and interactome (yellow). Metabolites are depicted as blue lines in the metabolome network and metabolites are shown in gray 3D chemical structures. GPCR structures in network are shown in yellow cartoon. **b** An overall workflow of ML-based structural pharmacology approach. Molecular docking was performed to generate 3D interaction features. Metabolites (gray sticks) derived from microbiota (blue cartoon) docked into pocket (blue funnel) of GPCR (cartoon cylinder); Structures of GPCR (multicolored 3D structures) were modeled by using AlphaFold2 based on multiple sequence alignment information; Then, diverse 3D interactions features, such as stability and ΔE of ligands in docking (red arrow), SASA (ligand mesh and protein gray surface), polar (yellow line) and hydrophobic (green line) interactions, were input for training ML models. The top-one GPCR ML model is prioritized as the Extra Trees regressor model (Tree diagram). Via comparing predicted GPCR-ML score in this model, we re-ranked the metabolites (yellow sticks) against GPCRs (gray cartoon) and selected the top-ranked metabolites for each GPCR.

### Multi-omics analysis reveals potential GPCR targets for Alzheimer’s disease

First, we examined the likely causal relationships between the GPCRome and AD by using Mendelian randomization (MR) analysis. Based on 3 large genome-wide association studies (GWAS) in AD^29–31^, 274 of 409 GPCRs were covered in our MR analysis. Among the 274 investigated GPCRs, 7 GPCRs were determined to be significantly related to AD (Fig. 2a). Of those, elevated expression of adhesion GPCR D1 (ADGRD1), GPR27, and vasoactive intestinal peptide receptor 2 (VIPR2) are associated with an increased risk of AD. By contrast, elevated expression of taste 2 receptor member 60 (TAS2R60), formyl peptide receptor 1 (FPR1), galanin receptor 1 (GALR1), and opsin 4 (OPN4) are associated with a reduced risk of AD. Specifically, bitter receptor TAS2R60 is associated with a substantially reduced risk of late onset AD (beta MR = −0.34, 95% confidence interval [CI] = −0.48 ± 0.20, FDR = 4.60 × 10^-4^). Three of the GPCRs identified in this exercise are also orphan GPCRs: GPR27, adhesion receptor ADGRD1, and photoreceptor OPN4. Expression of ADGRD1 has been reported in immune cells, and GPR27 is specifically expressed in the brain and is involved in neuronal plasticity and cognition^32–34^. For reference, the AlphaFold2 structural models of GPR27 and ADGRD1 are shown in Fig. 2b.

**Fig. 2.**
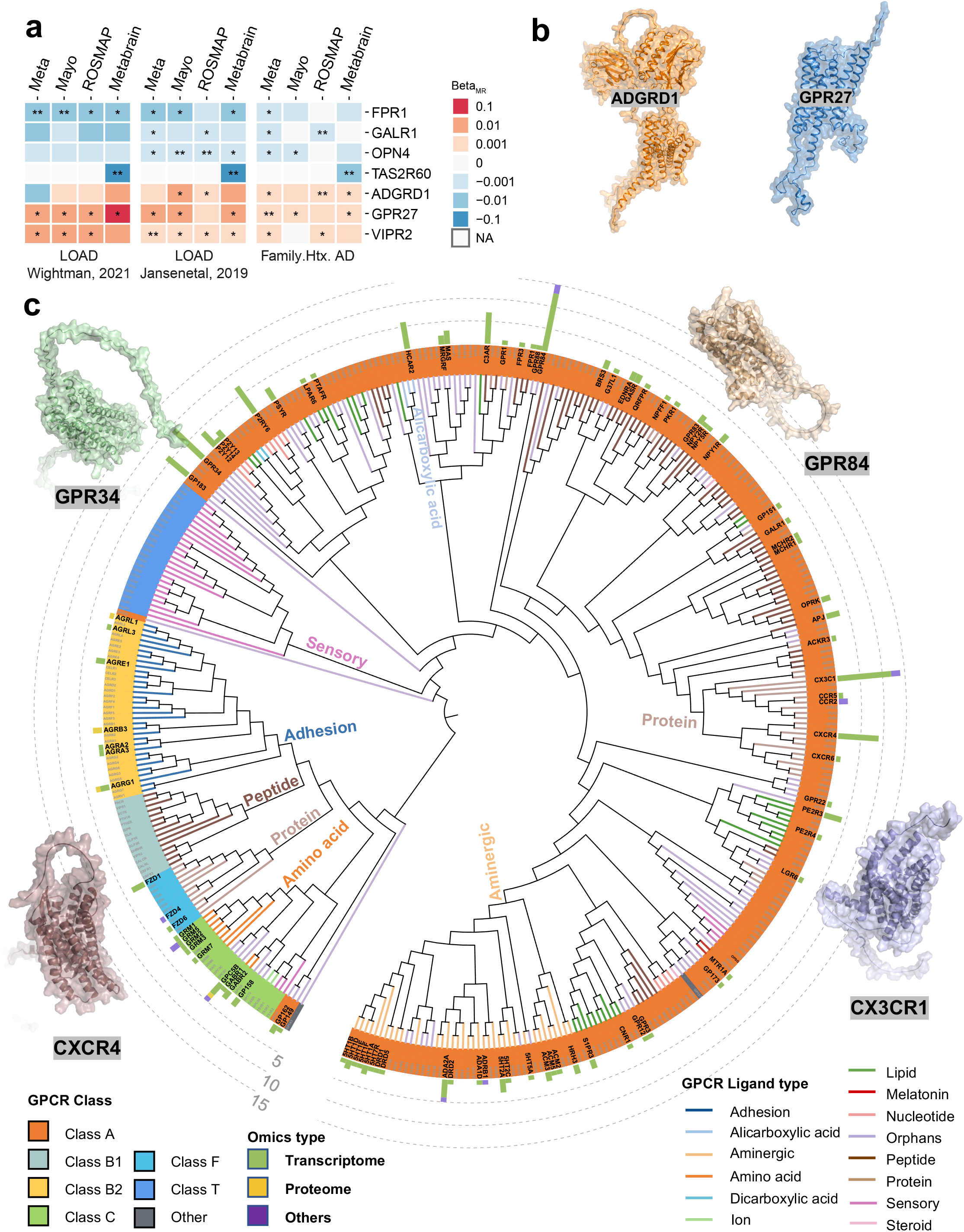
Multi-omics analysis reveals potential therapeutic targets for Alzheimer’s disease (AD) within the GPCRome. **a** AD associated GPCR targets suggested by Mendelian randomization (MR) analysis. 4 genomic datasets (Meta, Mayo, ROSMAP and Metabrain) in brain cortex region were inspected. Beta value > 0 indicates elevated expression level of GPCRs increase the likelihood of AD; Beta value < 0 elevated expression level of GPCRs reduce the likelihood of AD. The unit of beta is standard-deviation change in AD per 1-standard-deviation increase in gene expression. **denotes the significant cutoff using adjusted p value (FDR) < 0.05. * denotes significant *p* value < 0.05. **b** Protein models of top two prioritized AD-related orphan GPCRs, ADGRD1 and GPR27. AlphaFold2 structure models of both are depicted in orange and blue cartoon with surface, respectively. **c** Phylogenetic tree showing protein targets with AD multi-omics evidence across the GPCRome. Each branch represents one GPCR, labeled by UniProt protein name. The GPCRome is classified and indicated in colors of the outer circle. The ligand types of GPCRs are displayed on the out rim of the diagram. The number of multi-omics evidence is exhibited on the outer stacked bar chart, including Transcriptome (green), Proteome (yellow) and others (purple). GPCRs with at least one differential expression (DE) evidence are shown in black text on the tree rim. The outer gray dash line indicates the number of DE. 3D AlphaFold2 structure models of four GPCRs (GPR84, GPR34, CXCR4 and CX3C1) with the greatest number of DE evidences are depicted in cartoon with surface.

We next investigated the GPCRs for association with AD by analyzing bulk and single-cell transcriptomic or proteomic data from brain samples. Using 88 transcriptomic and proteomic datasets that we had compiled previously^35, 36^, we examined differential expression (DE) of 406 GPCRs between pathologic groups (e.g., AD) and control groups (e.g., cognitive healthy control). In total, 393 GPCRs were clustered to a circular phylogenetic tree based on protein sequence conservation (Fig. 2c). GPCR classes and the number of omics (i.e., number of datasets in which GPCRs were differentially expressed) are illustrated in Fig. 2c. In total, 92 GPCRs are suggested to be AD-associated (DE in at least one dataset), of which 23 differentially expressed GPCRs are orphan GPCRs. We then prioritized AD-associated GPCRs based on the degree of DE and identified the top 10 GPCRs with DE in at least 5 datasets (Fig. 2c), including GPR84, CX3CR1, GPR34, CXCR4, P2RY6, C3AR, GPR183, HCAR2, ADRA2A and GPRC5B. Among the top 10 differentially expressed GPCRs, GPR84, GPR34, GPR183 and GPRC5B are orphan GPCRs. Among all differentially expressed GPCRs, GPR84 is the strongest candidate, followed by C-X3-C motif chemokine receptor 1 (CX3CR1), C-X-C motif chemokine receptor 4 (CXCR4) and GPR34. GPR84 has been identified as a target of medium-chain fatty acids via mediating neuroinflammation^11^. Two chemokine receptors, CX3CR1 and CXCR4, also function to decrease neuroinflammation and regulate microglial activation^37, 38^. GPR34, another orphan GPCR, has been found to specifically expressed by microglia in AD^39^. Taken together, these genetic and multi-omics analyses reveal potential functional roles of the GPCRome in AD pathobiology, particularly for orphan GPCRs, including GPR84 and GPR34.

### Machine learning-based discovery of the gut metabolite-GPCRome

We next sought to identify gut metabolite-GPCRome interactions by combining AlphaFold2 and machine learning-based docking approaches (Fig. 1b). Here, we collected 516 unique, structurally characterized metabolites from a high throughput gut metabolomics study^40^. To identify potential GPCRome targets, we re-constructed 3D structures for 408 non-olfactory GPCRs using AlphaFold2 models. We observed that AlphaFold2-predicted GPCR structures containing high-confidence transmembrane helix (TM) regions (**Supplementary Fig. 1a**)^12^. We further compared the AlphaFold2-predicted and experimental structures deposited in Protein Data Bank (PDB) of 10 randomly selected GPCRs (**Supplementary Fig. 1b**). We found that the TM root-mean-square deviation (TM-RMSD) of selected GPCRs was less than 1Å, revealing the high-quality of AlphaFold2-predicted GPCR models. For pair comparison, refined crystal/Cryo-EM structures deposited in PDB (95 GPCRs with 482 structures, termed PDB structures) and homology models (404 GPCRs with 1031 structures) were also collected from GPCRdb^41^. In total, PDB structures and homology models revealed 77 identical GPCRs to 416 AlphaFold2-predicted models (408 GPCRs). Druggable pockets of GPCR structures (including PDB structures, AlphaFold2, and homology models) were characterized using Fpocket 2.0^42^. In total, we evaluated 1,680,096 metabolite-GPCR pairs connecting 408 GPCRs and 516 gut metabolites through molecular docking. We found a high Pearson’s correlation coefficient (R) of docking scores between 77 identical GPCRs shared in AlphaFold2 and experimental structures (R = 0.87, *p* < 2.2 x 10^-16^) (**Supplementary Fig. 2**), further revealing valid structures predicted by AlphaFold2 for metabolite docking studies.

We next developed a ML model to improve docking performance. Specifically, we assembled experimentally determined ligand-GPCR pairs with known binding affinity from GPCRdb^41^ and GLASS databases^43^. To implement the application domain of ML models between active GPCR ligands and gut metabolites, we focused on bioactive GPCR ligands having similar physiochemical properties to well-characterized human metabolites^44^ (**Supplementary Fig. 3**). In total, we harnessed 60,356 known ligand-GPCR pairs connecting 155 GPCRs and 38,117 bioactive ligands with experimental binding affinities (inhibition constant/potency, K_i_ value) serving as a training set. Then, 3D structural features from the docked ligand-GPCR complex were calculated to create ML models.

Among 11 evaluated ML algorithms, the Extra Trees Regressor model (R = 0.60) outperformed other ML approaches, such as Extreme Gradient Boosting (R = 0.50) and K neighbors (R = 0.43) (**Supplementary Table 1**). On an external test dataset, Extra Trees models (termed GPCR-ML score) also achieved similar performance (R = 0.63, *p < 2.2* x 10^-16^, Fig. 3a). By contrast, the docking score of external test dataset displayed weak correlation with binding affinity (R = 0.0055, *p* = 0.69) (**Supplementary Fig. 4**). To test over-fitting, we performed 10-fold cross-validation and found consistent performance (**Supplementary Table 2**). These observations suggest that the Extra Trees model derived GPCR-ML score enables us to identify the potency of metabolites against the GPCRs.

**Fig. 3.**
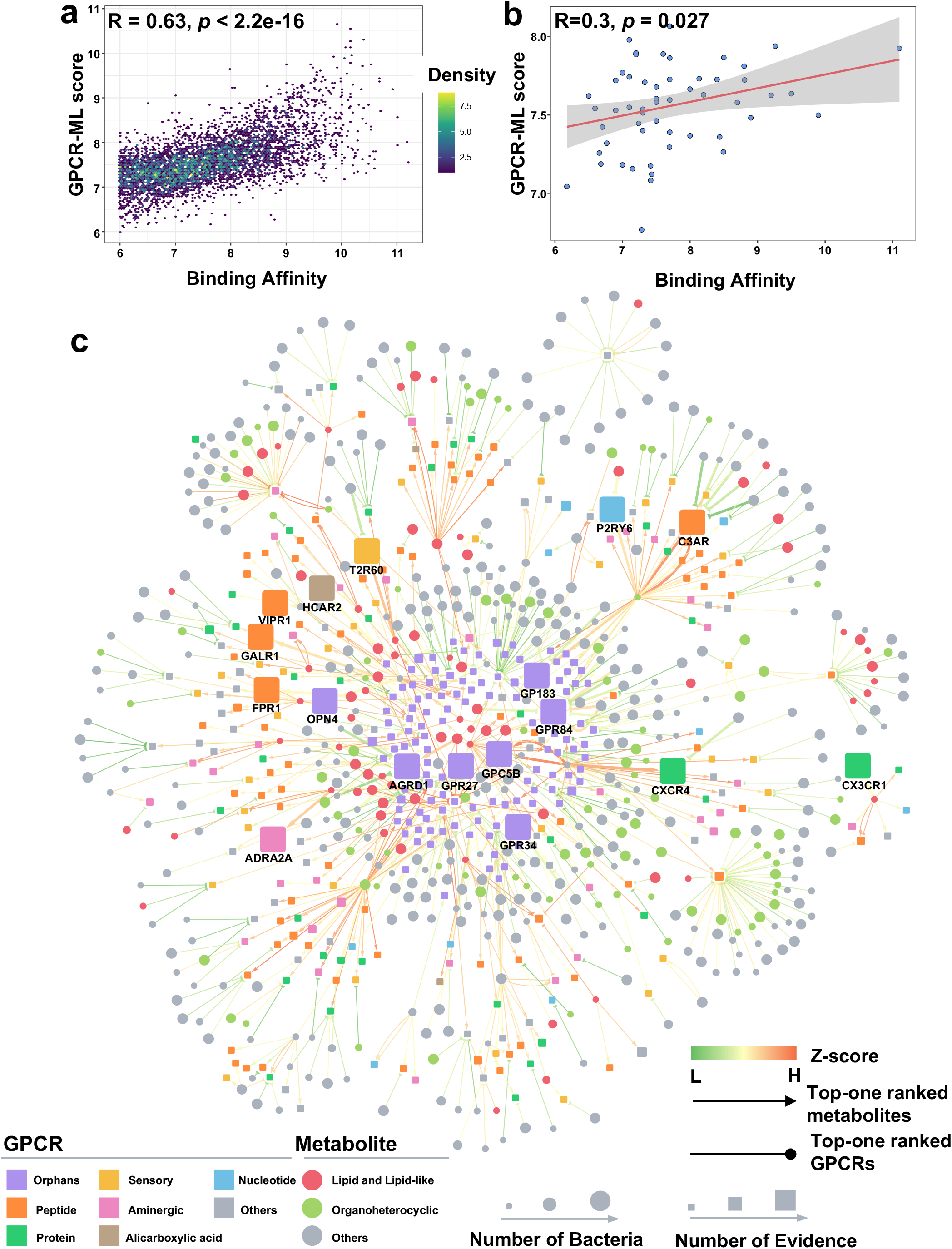
Machine learning-based discovery of the interactome of the gut metabolite-GPCRome network. **a** Performance of Extra Trees model on external test dataset. Binding affinity and GPCR-ML-score of 5,445 unseen metabolite-GPCR pairs were scatter plotted by point density. Pearson’s correlation coefficient R and *p* value are labeled. **b** Performance of Extra Trees model on a benchmark dataset. Binding affinity and GPCR-ML-score of 56 reported metabolite-GPCR pairs are scattered plotted in color blue. Regression of data was predicted by Linear Regression function, displayed as red line along with SD error (gray background). Pearson’s correlation coefficient R and *p* value are labeled. **c** An integrated network illustrates the bacteria-derived metabolite-GPCRome interactome. The top-one ranked metabolite of 369 GPCRs or the top-one predicted GPCRs of 516 metabolites were connected by a GPCR-ML score. Metabolite and GPCR are depicted as rectangle or circle nodes, respectively. GPCR-ML score (edge) was normalized by z-normalization and shown in green-orange color range based on size. The arrow edge means the top-one ranked GPCR target of metabolite, and the half-circle arrow line means the top-one ranked metabolites of GPCR. Hierarchical class of GPCR and chemical class of metabolites are indicated with different colors. The size of GPCR node is proportional to the number of MR and multi-omics evidence, while the size of metabolite node is proportional to the number of bacteria strains with higher metabolite abundance (abundance with | Log_2_FC | >=2 was shown).

To further validate the performance of ML models, we next conducted a benchmark study on an experimental metabolite-GPCR dataset. Previous studies have revealed that metabolites act as signaling molecules to activate GPCR pathways^4, 7^. Therefore, a set of 56 diverse metabolite-GPCR pairs against 34 GPCRs with agonistic activity (half-maximum effective concentration, pEC_50_ >= 6) were retrieved from GPCRdb together with the reported experimental data^7^. Of note, biological activities of all 46 metabolites (in 56 pairs) against GPCRs in the benchmark were determined by a β-Arrestin assay^7^. In addition to well-known GPCRs, recent understudied GPCRs, such as GPR183^4^, were also included, to improve the diversity of GPCR targets in the benchmark. We found that our GPCR-ML score achieved the best performance (R = 0.3, *p* = 0.027) compared to traditional AutoDock-derived docking score (R = 0.03, *p* = 0.82, Fig. 3b, **Supplementary Fig. 5**).

We further inspected 34 unique experimentally reported metabolite-GPCR pairs to evaluate the predictive performance of GPCR-ML score. We defined the top 10% of metabolites prioritized by GPCR-ML score as potential metabolites. As indicated in **Supplementary Fig. 6**, the hit rate at the top 10% level reaches 47.1% (16/34), revealing a ∼5-fold hit rate compared to traditional experimental screening (9%)^7^. Interaction between the bioactive metabolite serotonin and the 5HT1F receptor was predicted with the strongest GPCR-ML score. The reported bioactive metabolite 7,27-dihydroxycholesterol also rank as a strong potential metabolite pair for the orphan receptor GPR183, suggesting the potential power of this approach to rank metabolites for orphan GPCRs as well. We further evaluated the absolute error between GPCR-ML score and experimental binding affinity of all metabolite-GPCR pairs and found that 82.4% of the pairs (28/34) have a small absolute error (|absolute error| <= 1) (**Supplementary Fig. 6**). Taken together, our results indicate that the GPCR-ML score from the Extra Trees Regressor model offers a potential approach to identify metabolite-GPCR interactions, including orphan GPCRs.

### Microbiota-derived metabolite-GPCRome interactome

To inspect the interactions between microbiota-derived metabolites and the GPCRome, we predicted GPCR-ML scores of all metabolite-GPCR pairs (515 metabolites against 369 GPCRs) (*cf.* Methods). Metabolites were classified into 9 classes based on chemical structure similarity^40^. Among these classes, organic acids and derivatives (30.1%), organoheterocyclic compounds (18.3%), and lipids and lipid-like molecules (14.6%) were the top three categories (**Supplementary Fig. 7a**). The entire GPCRome was classified into 13 classes based on endogenous ligand type^45^ (**Supplementary Fig. 7b**). The largest class was found to be orphan GPCRs (33.6%), including Class A orphans (25.2%), Class B2 orphans (6.5%), and Class C orphans (1.9%). In addition, aminergic receptors composed 9.2% (34) of the 369 studied within the GPCRome. Aminergic receptors have been previously reported to be activated by gut commensals, which are microbes that reside on either the body surface or at the mucosa without normally harming human health^6^. To compare metabolite performance against each GPCR, the GPCR-ML scores of all metabolites were normalized to the maximum value (termed normalized score). For each metabolite, the mean of normalized scores across all GPCRs was calculated (**Supplementary Fig. 8**). Overall, the metabolite-GPCRome interaction network was divided into three groups based on hierarchical clusters of metabolites (**Supplementary Fig. 8a**): organic acid, lipid and lipid-like, and organic acid-benzenoids. These groups were based on their major metabolite type distribution within each group. Inter-groups displayed significantly distinct normalized score profiles (means of normalized scores over three groups are 0.43, 0.65 and 0.23, respectively; *p* < 2.22 x 10^-16^, Mann-Whitney U test, **Supplementary Fig. 8b**). Notably, the mean of the normalized scores of the lipid and lipid-like group was ∼1.5-fold of the organic acid group and ∼3-fold of the organic acid-benzenoids group. These observations reveal that GPCRs are more likely to be regulated by lipid and lipid-like molecules in comparison to other types of metabolites (**Supplementary Fig. 7a**).

Next, we sought to clarify the percentage of potential metabolites (prioritized by the top 10% of metabolites ranked by normalized scores) by testing each type of GPCR. To avoid a literature bias towards well-studied GPCRs and metabolites, we evaluated the percentage of potential metabolites by scaling the number of GPCR pockets and metabolites. The distribution of potential pockets displayed a similar pattern to that of GPCR types (**Supplementary Fig. 7b**), indicating that almost all GPCR classes (except for ion GPCRs) preferentially interact with lipid and lipid-like molecules (average percentage of potential metabolites is 19.4%, **Supplementary Fig. 9**), whereas GPCRs interact with organic acids and derivatives st a relatively lower percentage (average percentage is 2.4%). These results are consistent with our above metabolite-GPCR interaction network analysis (**Supplementary Fig. 8**).

We next focused on metabolites with the strongest GPCR-ML scores. To obtain a comprehensive network, we included the associations of top-one ranked GPCRs for each metabolite. The overall connectivity of the metabolite-GPCR network contains 884 nodes (369 GPCRs and 515 metabolites) and 884 edges (884 predicted metabolite-GPCR pairs, Fig. 3c). Among them, 4 pairs overlapped with experimentally reported metabolite-GPCR pairs (**Supplementary Fig. 10, Supplementary Table 3**). For GPCRome associations, the distributions of pairs by GPCR classes differed in 9 metabolite types (*p* = 0.028, *x*^2^ test). We evaluated the average number of pairs per GPCR. The top three GPCR types were melatonin receptors (3.5, 7/2), aminergic receptors (2.7, 92/34) and orphan GPCRs (2.7, 333/124) (**Supplementary Fig. 11a**). Of these, orphan GPCRs have the strongest connectivity with metabolites (333/884, 37.7%, **Supplementary Fig. 11b**). In this connectivity, there are 209 pairs in which metabolites interact with their top-one ranked orphan GPCRs, revealing that 40.6% (209/515) of metabolites prefer binding to orphan GPCRs. Over the whole associations of orphan GPCRs, lipid and lipid-like molecules are significantly associated with orphan GPCRs (75/333, 22.5%, *p* < 1 x 10^-4^, Fisher’s exact test). While 80.6% orphan GPCRs (100/124) have less than 3 pairs, ∼10 orphan GPCRs are still predicted to be targeted by over 5 metabolites, such as GPR12 and GPR83 (**Supplementary Fig. 12**). Two other major GPCRs involved in interactions with metabolites are peptide and aminergic receptors (21.5% and 10.4%, respectively). Previous chemical genetic screening indicated that aminergic GPCRs, such as histamine receptors and dopamine receptors, can be activated by gut-microbiota metabolites^6^. Here, lipids and lipid-like molecules were also observed as the largest categories in terms of pairings within the GPCRome (234/884, 26.5%) (Fig. 3c, **Supplementary Fig. 11b**), despite only comprising 14.6% of the entire metabolite datasets (**Supplementary Fig. 7a**). Orphan GPCRs were also found preferentially associate with lipid and lipid-like molecules (75/234, 32.1%, *p* < 0.05, Fisher’s exact test). These results are consistent with previous studies in which orphan GPCRs have been identified as sensors of lipid compounds, such as short-chain or medium-chain fatty acids^8, 11^. In addition, 34 metabolite-GPCR pairs consisted of the metabolite with strongest GPCR-ML score and its top-one ranked GPCRs (double edges in one pair shown in Fig. 3c), suggesting potential involvement in GPCR-metabolite associations. For example, the serotonergic GPCRs, 5HT2C and 5HT4R, have been suggested to be responsive to a variety of bacterial products^7^. We conclude that regardless of associations (whole interactions, top 10% metabolites, and top-one ranked metabolites/GPCRs), GPCRs preferably interact with lipid and lipid-like molecules, especially orphan GPCRs.

### Allosteric regulation of metabolites on the dark GPCRome

To characterize the ensemble of all discernible pockets, which is known as the -pocketome, of all metabolite-GPCR pairs, we next investigated pocket profiles in PDB structures, AlphaFold2-predicted, and homology models. Here, we defined orthosteric sites according to the endogenous ligand position and near key residues in experimental structures, while other pockets remote from the orthosteric site were regarded as the allosteric sites. In 77 identical GPCRs, the greatest occupancy of orthosteric sites is seen in PDB structures (61.4%), while occupancy in AlphaFold2-predicted and homology models are comparably lower (55.59% and 56.79%, respectively) (Fig. 4a). On the other hand, the pocket distributions of metabolite-GPCR pairs exhibited distinct patterns across three structural models (**Supplementary Fig.13**), revealing that AlphaFold2 may provide diverse binding sites compared to the PDB structures and homology models.

**Fig. 4.**
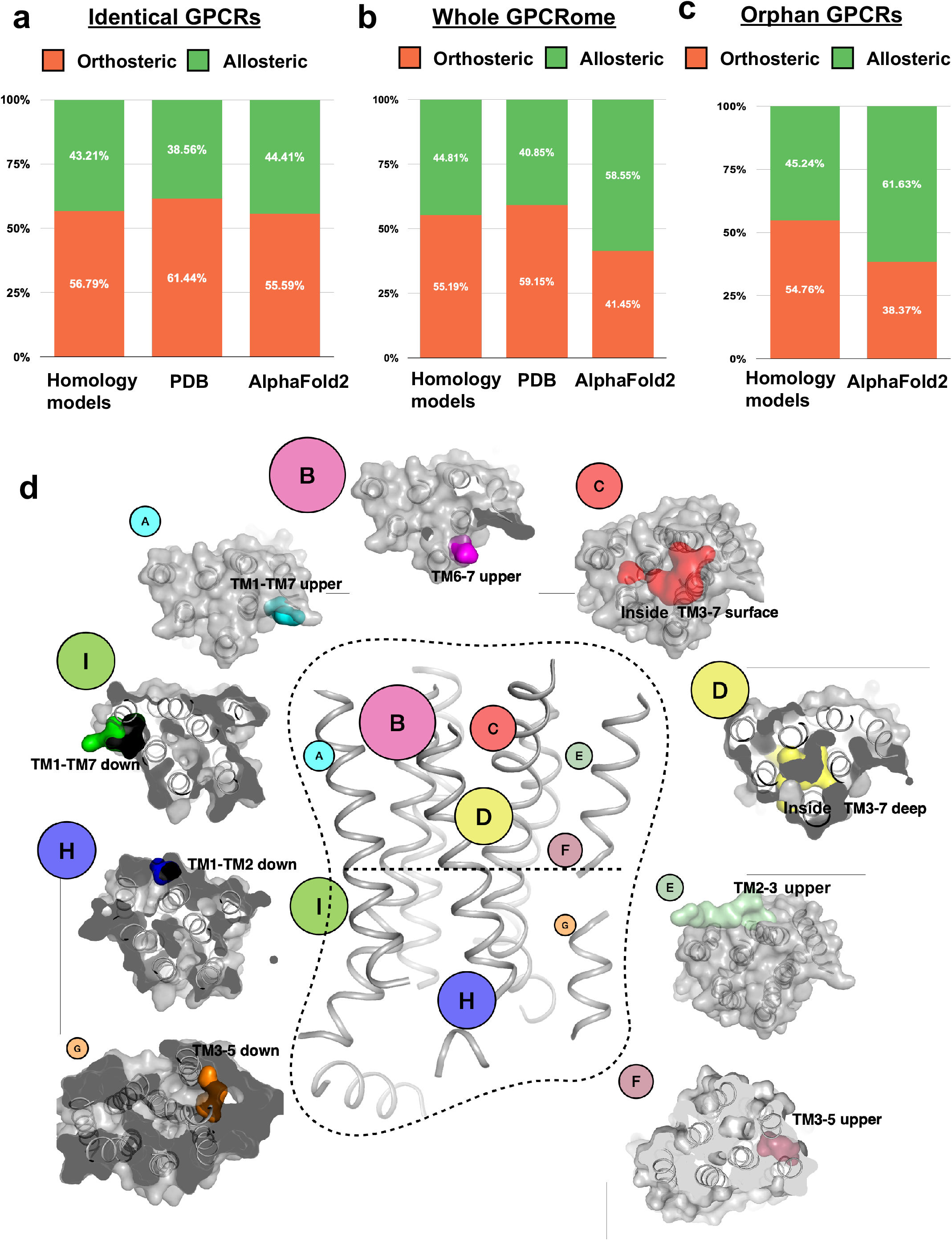
Allosteric regulation of metabolites on dark GPCRome. **a** A chart showing the binding pocket distribution of metabolites across 77 identical GPCRs shared in AlphaFold2-predicted models, PDB structures and homology models. The ratio of orthosteric pocket (orange) and allosteric pocket (green) are depicted. **b** A chart showing the binding pocket distribution of metabolite across all GPCRs in AlphaFold2-predicted, PDB structures and homology models, containing 369 AlphaFold2-predicted models, 92 PDB structures and 404 GPCRdb models. The ratio of orthosteric and allosteric pocket are depicted in color orange and green, respectively. **c** A chart showing the binding pocket distribution of metabolite across 99 orphan GPCRs in AlphaFold2-predicted and homology models. The ratio of orthosteric and allosteric pocket are depicted in color orange and green, respectively. **d** Pocket landscape of metabolite identified in Class C orphan GPCR models. Pockets of 3096 metabolite-GPCR pairs are clustered into 9 subtypes (A-I) of pocket based on their locations, and their position is shown on the GPCR model with multi-colored surface. Label size is proportional to the percentage of occupancy of metabolites. The AlphaFold2 model GRM5 is exhibited as a representative structure in the middle.

Since the above analysis mainly focuses on non-orphan GPCRs (98.7% of 77 identical GPCRs, **Supplementary Fig. 14**), we next analyzed the pocket profiles of the entire GPCR dataset, including orphan GPCRs. AlphaFold2-predicted models were shown to have a much larger occupancy of allosteric sites (58.55%) compared to the PDB structures and homology models (44.81% and 40.85%) (Fig. 4b). By contrast, the occupancy of allosteric site in AlphaFold2-predicted models was smaller than in PDB structures across 77 identical GPCRs (Fig. 4a). These results indicate that gut metabolites interact with orphan GPCRs largely via allosteric regulation. To avoid the bias introduced by different number of GPCRs contained within structural models, we compared the pocket profiles of 99 identical orphan GPCRs shared by AlphaFold2-predicted models and homology models. 61.63% of the pockets from AlphaFold2-predicted models are allosteric sites, which is considerably greater than that from homology models (45.24%) (Fig. 4c). Altogether, these observations suggest that gut metabolites are more likely to engage in allosteric binding with orphan GPCRs, which is consistent with recent experimental findings^46, 47^.

Allosteric modulators of Class C GPCRs have also been reported as potential candidates for treating AD^48^. Thus, we conducted pocket analysis of Class C orphan GPCRs. In total, 3,096 metabolite-GPCR pairs were investigated and classified into 9 pocket types (Pocket A-H, Fig. 4d), which were defined by their locations. Since the orthosteric site of Class C receptors is located in the extracellular domain, all 9 identified pockets are indicated as allosteric sites. Among them, we found one potential allosteric pocket previously known from crystal structures of Class C GPCRs (PDB: 4OR2, 6FFH), and 4 additional pockets that had also been reported in other GPCRs (PDB: 4XNV, 4MQT, 5NDZ and 5O9H). In addition, 4 untargeted allosteric sites were characterized from potential promising interactions. All of the above sites are in the lipid-receptor interface (Fig. 4d). Taken together, these potential allosteric modalities may not only facilitate the discovery of ligands for understudied GPCRs, but also have important implication in identifying the interaction of disease-relevant metabolites with orphan GPCRs allostery.

### The landscape of metabolite-GPCR associations in Alzheimer’s disease

A total of 293 pairs of AD-related GPCRs were mapped in the metabolite-GPCR interactome (containing 7 MR associated GPCRs and 88 multi-omics associated GPCRs) (Fig. 3c and Fig. 5a), accounting for 33.1% of all interactome associations (293/884). Among these pairs, orphan GPCRs formed the largest proportion (84/293, 28.7%) (Fig. 5a). The top three orphan GPCRs with the greatest percentage of associations were GPR12, GPR83 and GPR84 (Fig. 6a, **Supplementary Fig. 12**). Two other top-ranked GPCRs were peptide (71/293, 24.2%) and aminergic receptors (62/293, 21.2%), respectively. With respect to associations with orphan GPCRs, lipid and lipid-like molecules had the greatest percentage of metabolites (20/84, 23.8%). These results are in line with our interaction network analysis of top-one ranked metabolites and GPCRs associations (Fig. 3c).

**Fig. 5.**
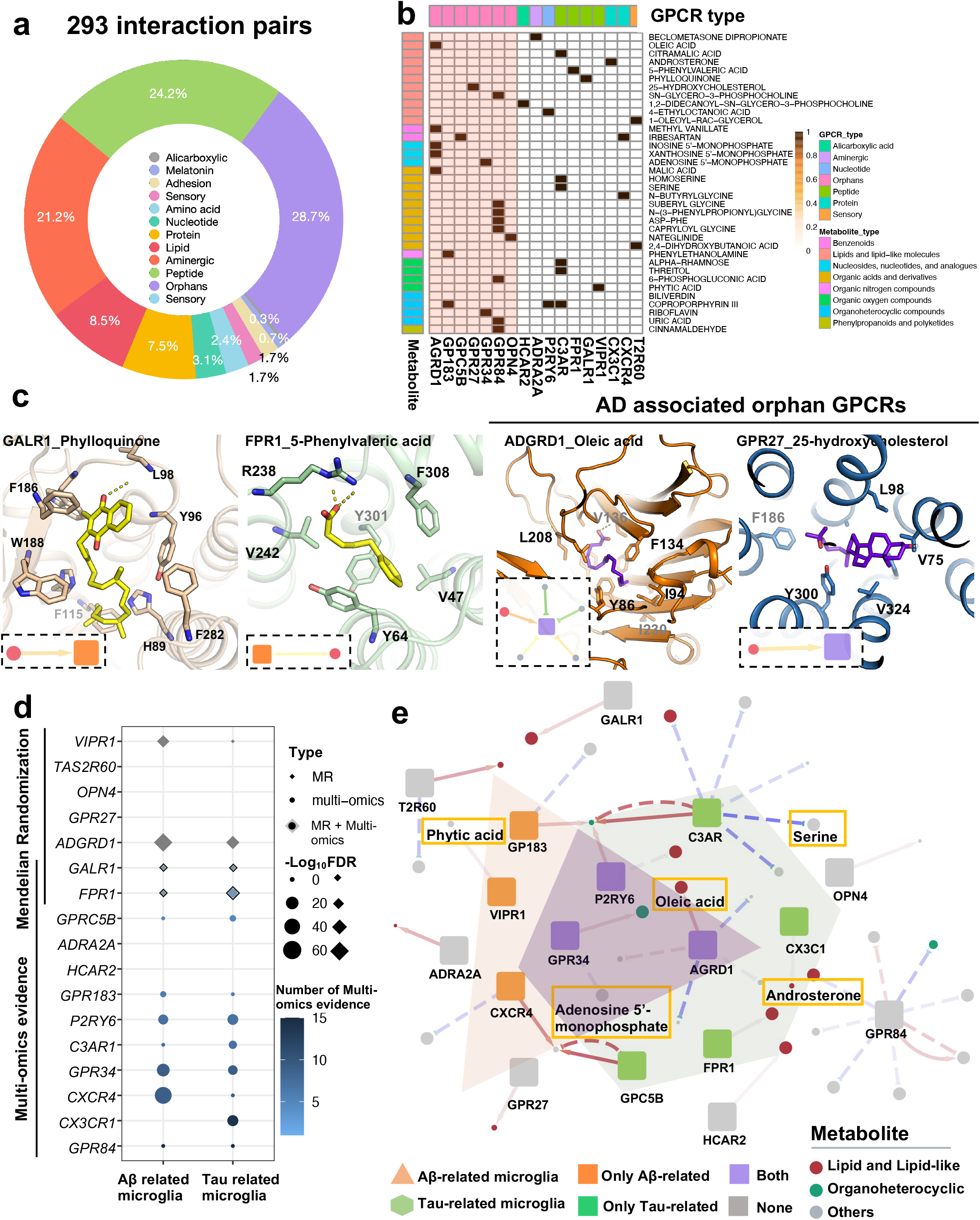
A landscape of microbiota-GPCR associations in Alzheimer’s disease (AD). **a** Percentage of metabolite-GPCR associations in AD. A total of 293 interaction pairs are classified by the ligand type of AD-related GPCRs. **b** Heatmap depicting associations of potential AD-related GPCRs with Mendelian randomization (MR) or multi-omics evidence. 41 selected metabolite-GPCR pairs are depicted in brown. The red columns display associations with orphan GPCRs. The GPCR-ML score is normalized to 0-1 by adopting Min-Max normalization. **c** Binding modes of four AD-related GPCRs with their top-one ranked metabolites. GALR and FPR1 are two GPCRs possessing both genetic and transcriptomic evidence; ADGRD1 and GPR27 are two MR-suggested orphan GPCRs. The network of four GPCRs associated with metabolites were depicted. Metabolites paired with two non-orphan GPCRs are depicted in yellow sticks, while that of orphan GPCRs are shown in purple sticks, proteins are shown in cartoon with multi-color. Key residues around metabolites were exhibited in sticks. **d** Differentially expressed GPCRs in Aβ related microglia or tau related microglia. The significance of differential expressed genes were represented by −Log_10_FDR value and indicated by size of circle. FDR < 0.05, indicates significant different in disease-associated microglia (DAM) (vs. homeostasis-associated microglia (HAM)). The number of multi-omics evidence are also displayed in color white-blue. **e** An overall landscape of metabolite-GPCR associations in AD and AD-associated microglia. 41 pairs between GPCRs (7 MR-related GPCRs and 10 muti-omics related GPCRs), and 35 metabolites were shown in the network. The symbol of arrow or circle are the same as depicted in Fig. 3. Literature reported metabolites that are involved in neuroinflammation regulation in microglia are indicated by an orange frame.

**Fig. 6.**
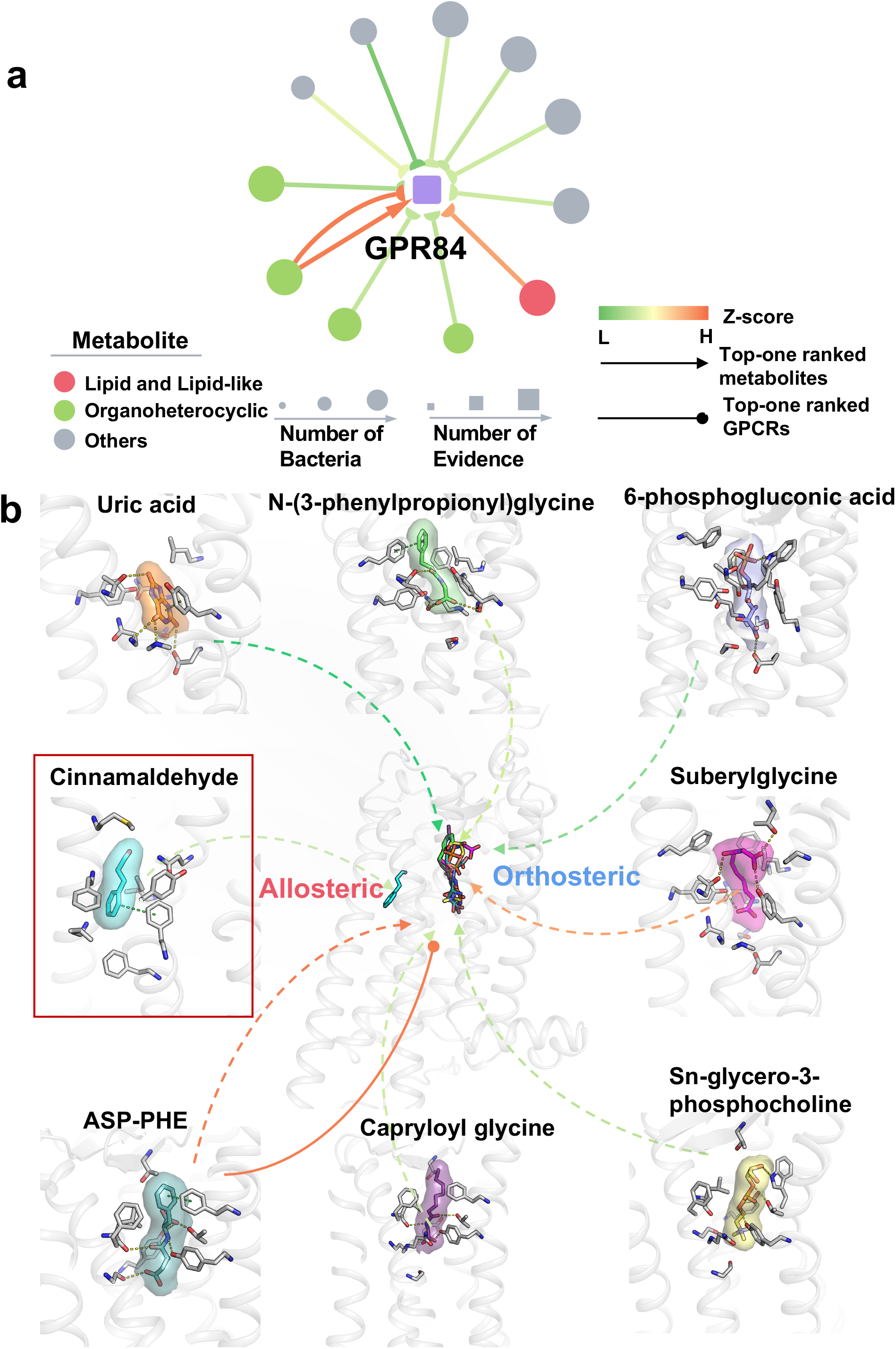
Pocket distribution of orphan GPR84. **a** GPR84-centered metabolite-GPCR network. Symbols shared with Fig. 3. **b** 3D interaction network on orphan GPR84. Metabolite is shown in sticks with surface and protein structure of GPR84 is shown in gray cartoon. Detailed binding modes of 8 metabolites are exhibited on the structure of GPR84, including 7 metabolites with their top-one ranked GPR84 and one top-one ranked metabolite from GPR84 docking results. Two pockets, including one orthosteric pocket and one allosteric pocket, are indicated. Key residues around metabolites are shown in gray sticks. The chemical name of metabolite is labeled in black. The arrow lines are displayed as same as the network shown in Fig. 3c.

Furthermore, when we exclusively considered only potential AD-associated GPCR with multi-omics evidence (10 GPCRs with DE in more than 5 datasets) and likely causal GPCRs with genetic evidence (7 MR associated GPCRs), 41 associations (with 35 metabolites) remained (Fig. 5b). Of these, more than a half (54%) were related to orphan GPCRs. As expected, most of the associated metabolites (11/41, 26.8%) were lipids and lipid-like molecules.

Among the GPCRs associated with AD, two GPCRs possessed both genetic and transcriptomic evidence, including galanin receptor 1 (GALR1) and formyl peptide receptor 1 (FPR1) (Fig. 2). Both GPCRs only interacted with one metabolite (Fig. 5b). The metabolites paired with GALR1 and FPR1, phylloquinone and 5-phenylvaleric acid respectively, were lipid-like molecules. Their association network and binding modes are depicted in Fig. 5c. FPR1 and GALR1 are both peptide receptors that are mainly expressed in the brain and gastrointestinal tract. FPR1 has been found to regulate mucosal immune systems, and GALR1 has been associated with cognitive decline in AD^49, 50^.

T2R60, the GPCR with the most MR evidence, interacts with two top prioritized metabolites, 1-oleoyl-rac-glycerol and 2,4-dihydroxybutanoic acid. Another two MR-supporting orphan GPCRs, ADGRD1 and GPR27 were also examined (structures shown in Fig. 2b). ADGRD1, was found to be bound by 5 metabolites, while GPR27 was found to be bound by only one metabolite (25-hydroxycholesterol) (Fig. 5c). Here, the predicted structure of ADGRD1 consists of a large N-terminus (Fig. 2b). Oleic acid, a top-one predicted metabolite of ADGRD1, binds to its allosteric domain. Notably, a previous study reported that ADGRD1 was agonized by binding to its N-terminal stalk^51^. 25-hydroxycholesterol displayed the strongest GPCR-ML score for GPR27, with binding to an extracellular site that is different from the reported binding site^32^.

Among all investigated GPCRs, GPR84 was identified as a candidate AD target for it having the strongest multi-omics evidence (Fig. 2c). In addition, GPR84 also possessed the greatest number of pairs with top-ranked candidate metabolites (8/36, 22.2%, Fig. 5b). Fig. 6 displays the association network and potential binding sites of GPR84. While only two potential pockets were characterized, one of them was revealed as an allosteric pocket. This allosteric pocket in the GPR84 model was observed to be hydrophobic, allowing it to interact with cinnamaldehyde, one of the examined metabolites (Fig. 6b). While each of most of the potential AD-associated GPCRs possesses 1-2 associations (Fig. 5b), complement receptor C3AR binds to 6 prioritized metabolites (**Supplementary Fig. 15a**). 4 out of the 6 are carboxylic acids and derivatives. Consistently, a few carboxylic acid compounds were reported to activate C3AR in vitro^52^. The C3AR-centered association network and binding mode of the top-one ranked metabolite coproporphyrin III is shown in **Supplementary Fig. 15b**. Coproporphyrin III structurally binds to an allosteric pocket located at the interface of the lipid and receptor, and has been suggested to induce expression of pro-inflammatory cytokines^53^.

### Metabolite-GPCR associations in AD-associated microglia

Because disease-associated microglia (DAM) have been previously shown to be pathologically associated with AD^54^, we further inspected the differential GPCR genes expressed in DAM by using single nucleus RNA-sequencing data analysis, including in Aβ and Tau related microglia (GSE148822)^55^. We found that 3 out of 7 MR recognized AD-associated GPCR genes (*FPR1*, *ADGRD1* and *VIPR1*), and 7 out of a top 10 multi-omics identified AD-associated GPCR genes (*CX3CR1*, *CXCR4*, *C3AR1*, *GPRC5B*, *GPR34*, *GPR183*, *P2RY6*), were differentially expressed in DAM compared to homeostasis-associated microglia (HAM) (FDR < 0.05) (Fig. 5d). In addition, *GPR34*, *P2RY6* and *ADGRD1* were differentially expressed in both Aβ-related microglia and Tau-related microglia. These results indicate that the above-mentioned GPCRs may be involved in AD through DAM. Next, we investigated the association of these GPCRs with metabolites (Fig. 5e). Among metabolites paired with the DAM-associated GPCRs (19/35), 5 metabolites have been reported to regulate neuroinflammation in microglia, including adenosine 5’-monophosphate (paired with GPR34)^56^, androsterone (paired with CX3CR1)^57^, oleic acid (paired with AGRD1)^58^, phytic acid (paired with VIPR1)^59^ and serine (paired with C3AR)^60^. Taken together, we showed proof-of-concept of metabolite-GPCR associations in a cell type-specific manner using AD-associated microglia as an example.

### Discovery of microbiota-derived metabolite-GPCR associations in Alzheimer’s disease

A previous study revealed that the abundance of *Eubacterium rectale* bacteria in patients with AD-related amyloidosis was significantly reduced compared to control subjects^61^. Therefore, we next examined which metabolites from *Eubacterium rectale* is significantly increased by mining its metabolite profiles and analyzing their reported abundance^40^ (**Supplementary Fig. 16**). Four metabolites (including indolelactic acid, 2’-deoxyuridine-5’-monophosphate, 2-(4-hydroxyphenyl)-propionic acid and dimethylbenzimidazole) were significantly elevated by at least four-fold (log_2_FC ≥2, relative fold change over the germ-free control) (Fig. 7a). Two of these metabolites (indolelactic acid and dimethylbenzimidazole) were found to be organoheterocyclic compounds involved in the tryptophan metabolic pathway and neurological regulation^62, 63^. Next, we explored the top 10 predicted GPCRs and mapped them to our MR and multi-omics analyses results. Six unique AD-associated GPCRs were identified, including apelin receptor (APJ), adrenoceptor Alpha 1D (ADRA1D), adrenoceptor beta 1 (ADRB1), prokineticin receptor 1 (PKR1), adhesion GPCR B3 (AGRB3), and lysophosphatidic acid receptor 6 (LPAR6) (Fig. 7b). In addition, three AD-related GPCRs (APJ receptor, ADRA1D and LPAR6) were predicted to be the targets of 2-(4-hydroxyphenyl)propionic acid. Notably, the APJ receptor is shared by three out of four metabolites and also predicted as the top-one ranked target (Fig. 7b). The APJ receptor-coding gene was found to be over-expressed in two human brain transcriptomic datasets, and have been suggested as a promising target in AD therapies^64^. Our results suggest bacterium *Eubacterium rectale* may shape AD in part by engaging the APJ receptor.

**Fig. 7.**
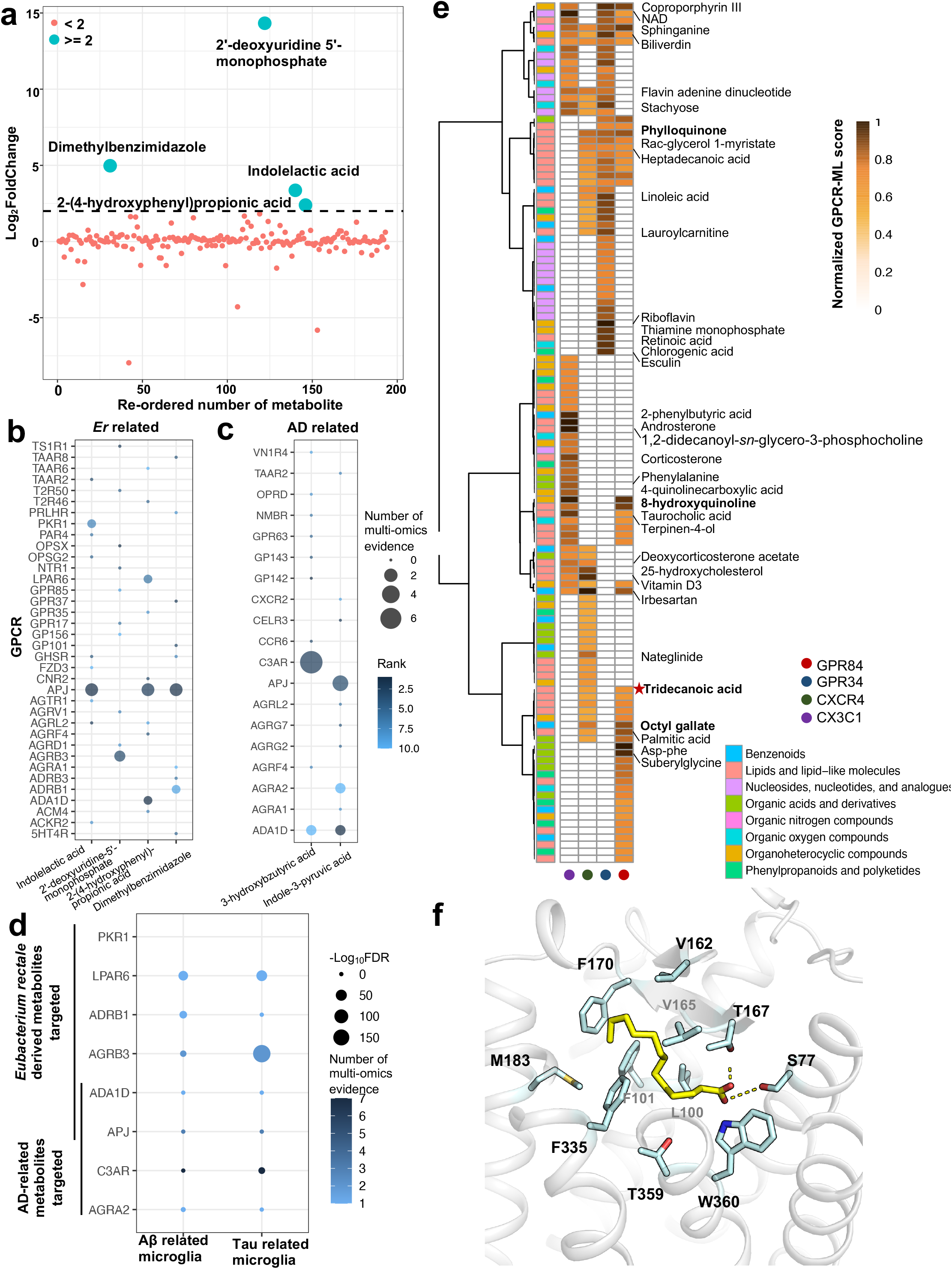
Discovery of microbiota-derived metabolite-GPCR associations in Alzheimer’s disease (AD). **a** Abundance of metabolites derived from *Eubacterium rectale.* 194 related metabolites are re-numbered, and their abundance are represented by using Log_2_FoldChange (Log_2_FC). The Log_2_FC of abundance that over than 2 are indicated with green circle, otherwise with red circle. The dash line means Log_2_FC equal to 2. **b** Bubble plot illustrating top 10 GPCR targets of metabolites derived from *Eubacterium rectale.* The size of bubble shows the number of multi-omics evidence of metabolite. The color range shows the docking ranks of identified GPCR according to the GPCR-ML-score. **c** Bubble plot illustrating top 10 GPCR targets of two reported AD-related metabolites. **d** Differential expressed genes of AD-associated GPCRs targeted by *Eubacterium rectale*-derived or AD-related metabolites in Aβ related microglia or tau related microglia. Six AD-associated GPCRs targeted by 4 *rectale*-derived metabolites and 4 AD-associated GPCRs targeted by two AD-related metabolites were inspected. The significance of differential expressed genes (FDR < 0.05) are represented by −Log_10_FDR value and indicated in size of circle. The number of multi-omics evidence were also displayed in color white-blue. **e** Heatmap depicting top 50 metabolites of 4 best AD-related GPCRs (CX3C1, CXCR4, GPR34 and GPR84) with strongest multi-omics evidence. The GPCR-ML score is normalized to 0-1 by adopting Max-Min normalization. Metabolites are hierarchically clustered (Ward’s D method) using Manhattan distance between the normalized GPCR-ML score across all taxonomies. The name of top 10 metabolites for each GPCR were exhibited. Important metabolites of GPR84 were addressed with bold label. The metabolite tridecanoic acid of GPR84 are labeled with red star. **f** Detailed binding mode of tridecanoic acid and orphan GPR84. Molecule is shown in yellow sticks and protein is shown in gray cartoon. Yellow dash line indicates hydrogen bond. Key residues near the binding pocket are displayed in cyan sticks.

Growing evidence suggested that two short-chain fatty acids, 3-hydroxybutyric acid and indole-3-pyruvic acid reduced AD related neuropathology^8, 65^. The MR and multi-omics analysis of the top 10 ranked targets were next inspected (Fig. 7c). Two AD-related GPCRs, ADRA1D and C3AR, were identified as the targets of 3-hydroxybutyric acid. C3AR is a well-defined immune modulator that is suggested to mediate neuroinflammation and Tau pathology in AD^66^. In our multi-omics analysis, C3AR is differentially expressed in 7 mouse transcriptomic datasets, including 3 microarray and 4 bulk RNA-seq datasets. Of those, C3AR is significantly upregulated in 3 datasets (FDR < 0.05, AD vs health control). For example, C3AR is found to be significantly overexpressed in APP/PSEN1 mouse model (Log_2_FC = 0.81, FDR = 7.16 × 10^-3^, 4 AD mice vs. 4 health control)^67^. In addition, microbiota-derived metabolite n-butyrate has been reported to regulate the gene expression of a subunit of C3AR^68^. Likewise, three multi-omics evidenced AD-related GPCRs were identified as the targets of metabolite indole-3-pyruvic acid, including ADRA1D (one transcriptomic evidence), adhesion receptor AGRA2 (one transcriptomic evidence) and APJ receptor (three transcriptomic evidence). We conclude that our strategies have the potential to identify the mechanism-of-action between metabolites and AD via specifically targeting GPCRs, e.g., APJ receptor and C3AR.

To test whether gene expression of potential AD-associated GPCRs targeted by above *Eubacterium rectale*-derived or AD-related metabolites were also altered in AD-associated microglia, we further performed a differential expression analysis between DAM and homeostasis-associated microglia (HAM) (Fig. 7d). Here, we found that three out of six genes of predicted AD-associated GPCRs targeted by *Eubacterium rectale*-derived metabolites were differentially expressed in DAM (FDR < 0.05, vs. HAM), including *ADRB1*, *ADGRB3* and *LPAR6*. The associated metabolites are dimethylbenzimidazole, 2’-deoxyuridine-5’-monophosphate and 2-(4-hydroxyphenyl)-propionic acid, respectively. Interestingly, dimethylbenzimidazole is a component of vitamin B12 that has been reported to suppress microglial activation^69^. Also, among 4 predicted AD-associated GPCR genes of two AD-derived metabolites, *C3AR1* (encoding protein C3AR, paired with 3-hydroxybutyric acid) was differentially expressed in DAM, which is in line with a previous study^66^. Notably, 3-hydroxybutyric acid has been reported to alleviate neuroinflammation in AD^70^, and we speculate that it may modulate AD pathology by acting on C3AR in microglia.

As potential AD-related targets were identified in our multi-omics data, we next asked whether we could identify potential bioactive metabolites by targeting these AD-associated GPCRs. To this end, we sought to analyze the metabolites of the four GPCRs with the greatest weight of multi-omics evidence for relation to AD: GPR84, CX3CR1, CXCR4 and GPR34 (Fig. 2c). The top 50 metabolites (top 10%, potential metabolites by our definition) were prioritized by normalized scores, and the chemical names of the top 10 metabolites are indicated in Fig. 7e. For example, we visually examined the potential metabolites of GPR84 since it is the strongest AD-associated GPCR in multi-omics analysis. 8-hydroxyquinoline, which is 3^rd^ ranked in metabolites associated with GPR84, and its derivatives have been reported to display significant inhibitory effects against Aβ aggregation in AD^71^, and here this metabolite also ranked 4^th^ among potential metabolites of CX3C1 (Fig. 7e). Another 7^th^ ranked potential metabolite, octyloctyl gallate, has been reported to markedly reduce cerebral amyloidosis in a mouse AD model^72^, and here it also ranked 6^th^ among potential metabolites of CXCR4. Another important potential metabolite, phylloquinone (vitamin K1), ranked 9^th^ for GPR84 and 5^th^ for CXCR4, and is highly expressed in the *Bacteroides* genus (log_2_FC > 10, **Supplementary Fig. 16**). Notably, previous studies have suggested a direct association between phylloquinone and cognitive function in AD patients^73, 74^. Overall, half of the potential metabolites are lipid and lipid-like molecules associated with GPR84 (25/50). Of those, 11 were fatty acids and derivatives, including two medium-chain fatty acids (suberylglycine and tridecanoic acid), agreeing with a previous report that medium-chain fatty acids activate GPR84^11^. The binding mode of suberylglycine with GPR84 is shown in Fig. 6b. Because tridecanoic acid (Fig. 7f) has previously shown an pEC50 affinity of 5.77 in the GPR84 β-arrestin assay^11^, we investigated its binding mode with GPR84 (Fig. 7f). The carboxyl group of tridecanoic acid forms two hydrogen bonds with residues Thr167 and Ser77, and its carbon chain points to a hydrophobic region that interacts with hydrophobic residues, such as Leu100, Phe101 and Phe170. Because our ML training dataset does not contain either GPR84 or tridecanoic acid, the strength of our GPCR-ML score is further underscored.

In summary, these findings suggest that potential application of our integrated network-based approaches to identify mechanism-of-action of AD-related metabolites may offer potential metabolite-based treatment approaches in AD.

## Discussion

Much effort in the field has been devoted to developing therapeutics based on targeting disease-modifying modalities or genetic factors in AD. However, thus far over 99% of clinical trials have failed^75^, highlighting the heterogenous etiopathology and pathogenesis of AD. Thus, there is an urgent need to uncover potential targets from a chemical perspective. The microbiome, a prominent member of the environmental exposome, has been suggested to mediate communication between the ecosystem of the gut and human brain in AD pathologies^17^. While recent advances indicate that alterations of bacteria or microbiota-derived metabolites modulate neurological disorders^76^, insufficient understanding of the potential targets of microbial products largely limits novel therapy development and clinical translation. Notably, microbiota-based therapeutics have been employed to treat autism spectrum disorder^77^, which serve as a proof of principle that other novel therapies for brain health, including AD, may also be derived by targeting microbiota dysbiosis. In our study, we developed a conceptual ML-based structural systems pharmacogenomics framework based on cheminformatics and bioinformatics data. This has enabled us to discover potential targets of microbiota-derived metabolites rapidly and effectively. Here, we concentrated on one of the key therapeutic targets in AD, the GPCRome. We clarified the interactome of large-scale metabolite-GPCR pairs, providing a landscape of AD-related GPCRs with genetic and multi-omics evidence, as well as their metabolite interaction partners. Furthermore, we demonstrated that metabolites may be allosteric regulators of GPCRs by preferential binding to allosteric sites in AlphaFold2 models, especially lipid and lipid-like molecules. Finally, we validated our approaches by investigating potential GPCR targets of bacteria-derived and AD-associated metabolites.

ML and artificial intelligence applications have been developed to predict the bioactivity of ligands binding to GPCRs^78^. Most computational methods are based on 1D or 2D features of ligands or proteins, such as multilayer perceptron (MLPs) and convolutional neural networks (CNNs), or machine-learning based scoring functions based on distance features or interatomic pairs^79, 80^. However, these methods are usually limited by training set quantity and easy-overfitting issues. Recently, one scoring function based on 3D fingerprints representing ligand-protein physical interactions was reported^81^. Importantly, this model outperformed other reported score functions, suggesting the reliability of the interaction features. To construct our metabolite specific GPCR-ML model, three strategies were adopted: (1) We implemented a high quality of ligand-GPCR training set of bioactive ligand-GPCR pairs with high bioactive potency (pKi >= 6); (2) We utilized a large quantity training set that are metabolite mimics. To mitigate the risk of this leading to unreliable data derived from property differences between bioactive ligand and metabolite datasets, we applied a physiochemical properties cutoff calculated from a large-scale metabolite dataset to improve the chemical similarity in training set; (3) We utilized high quality 3D features in our analysis. To improve the performance of docking, for example, we used high quality AlphaFold2 models and increased ligand sampling space in docking, since features are largely dependent on the docking pose. The Advanced Deep Learning based AlphaFold2 algorithm may provide potential reliable models for GPCRs, especially for unstructured orphan GPCRs. To validate the performance of AlphaFold2, we manually and individually inspected all structures of non-olfactory GPCR models, and observed that they all contained confidently predicted transmembrane helix (TM) regions^12^ (some of the models are displayed in **Supplementary Fig. 1a**). This substantiates the use of molecular docking with the AlphaFold2 models, since reported conserved pockets of GPCRs are mainly distributed on the TM domain^82^. We also randomly compared the structural differences of AlphaFold2 and crystal structures in 10 GPCRs (**Supplementary Fig. 1b**). The TM root-mean-square deviations (TM-RMSD) of all selected GPCRs are very low (less than 1 Å), pointing to the high-quality of the AlphaFold2-modeled GPCR structures. To avoid overfitting, we removed features with higher multicollinearity (threshold = 0.95). To improve the quality of the training set, ligand-GPCRs with lower affinities (pK_i_ < 6) were not considered in the training set. This protected us from overstating the predicted scores of poor binding metabolites, and enabled us to focus on the ranking of metabolites rather than their scores.

The quality of our predicted metabolites with GPCR-ML score is evident from two results. First, compared with a scoring function validated on the CASF-2016 dataset, GPCR-ML score has a comparable performance with X-Score^83^, yet lower than ΔVina-RF20^84^. The feature importance of Solvent accessible surface area (SASA) (**Supplementary Fig. 17**) is in accordance with previous reports^81^, while the ligand stability features in our calculation, including ligand RMSD and conformational energy differences (ΔE), are not the most top features, possibly due to different ML models rather than datasets. Second, compared with docking score, binding affinity has a good correlation with GPCR-ML score on the benchmark dataset, even though our bench dataset is of a relatively small quantity (Fig. 3b).

Importantly, we identified several potential AD-related GPCRs. A few of them, such as CX3C1, CXCR4 and C3AR, are chemokine receptors, activation of which have been suggested to have an important neuroprotective role in AD, such as C3AR for attenuating Tau pathology^85^. T2R60, a bitter taste receptor, has been suggested to be expressed in the gastrointestinal tract^86^, yet its functions in neurological disorders are under-investigated. Previous studies suggested that TAS2R receptors could be involved in immune regulation^87^, and we thus hypothesize that T2R60 may be a proof-of-concept AD target by playing a role in peripheral inflammation. GPR183, another immunomodulatory EBI2 receptor, has been recently reported to be agonized by the metabolite lauroyl tryptamine derived from *Eubacterium rectale*^88^. Furthermore, HCAR2 (GPR109A), a microglia receptor that was induced by amyloid pathology in AD^89^, was also identified as being agonized by the metabolite nicotinic acid derived from *R. guavus*^7^. A FDA-approved drug Niaspan was reported to activate HCAR2 to modulate amyloid pathology and neuronal dystrophy in AD^90^. Notably, the top-ranked metabolites of GP183 and C3AR are both coproporphyrin III. Even though our ML model has improved accuracy, more metabolites need to be inspected, for example the top 50, rather than just those topping the list. Our findings are consistent with a previous report that GPCRs may play an important role in AD pathology^91^, including amyloid, tau, inflammation and neurodegeneration, pointing towards pharmacologic approaches that target AD-associated GPCRs and providing potential insights into the identification of related biomarkers.

GPCRs mediate diverse pathway signaling by different ligands, making it important to analyze their pocket landscape and illustrate their molecular pharmacology^5^. So far, how distinct metabolites structurally regulate GPCRs remains largely unknown, particularly for unresolved orphan GPCRs^4, 92^. Since allosteric modulators have been advancing as drug candidates in treating CNS disorders that may reduce the risk of receptor oversensitization^93^, we investigated the occupancy of orthosteric and allosteric pockets in our metabolite-GPCR complex. Most metabolites in our dataset bind to allosteric pockets in AlphaFold2 orphan GPCR models (Fig. 4a-c), which is consistent with previous reports that metabolites may activate orphan GPCRs, serve as allosteric modulators, and play a role in allosteric regulation of CNS disorders^4, 94, 95^. Indeed, several allosteric modulators of Class A and Class C GPCRs have entered clinical trials for treating neurological disorders^93^. For example, the negative modulator of mGluR5, one receptor of Class C GPCRs, has shown promising effects in treating anxiety and Parkinson’s disease in phase II trials^96^.

Together, our findings highlight the utility of the systems pharmacogenomics framework and multi-omics approaches to uncover the molecular relationships between gut-microbiota metabolites and GPCRome targets in AD. We envision that our predicted communications would provide essential sights into future experimental or clinical validations on therapeutically regulations of gut microbiota for treating AD.

## Methods

### Datasets curation

833 metabolites derived in vivo and in vitro from 178 human gut bacteria strains were collected from recent studies conducted by S. Han et al.^40^. Duplicate chemicals were removed based on PubChem_ID and chemical name, resulting in 516 molecules (**Supplementary Data 1**). The metabolic profiles and abundance in diverse bacteria of these metabolites were re-analyzed. 15,092 metabolites were assembled from SMPDB database^97^ for molecular physiochemical property calculations. 1,045,681 entries from GPCRdb^98^ and 321,881 bioactive ligand-GPCR entries from GLASS^43^ databases were collected for the ML training set. 56 reported metabolite-GPCR pairs from GPCRdb^41^ and a previous study by D. A. Colosimo et al.^7^ were collected for the benchmark. 416 high quality structural models of 408 known human non-olfactory GPCRs were assembled from AlphaFold2 Protein Structure Database^12^. 95 GPCRs with 482 crystal structures and 404 GPCRs with 1031 homology models were collected from GPCRdb database. The number of identical GPCRs among three models is 77. The list of GPCRs is available in **Supplementary Data 2**.

### Molecular docking and interaction analysis

2D structures of molecules, including metabolites were converted by using Open Babel, and 3D structures were generated and prepared by using LigPrep module (Schrödinger Inc., version 2020.1). All protein structures, including AlphaFold2-predicted, PDB structures and homology models, were prepared by using the Protein Preparation Wizard module (Schrödinger Inc, version 2020.1). Also, molecules and proteins were prepared by using Autodock utilities^99^ for docking. To identify novel pockets as much as possible, Fpocket suite (version 2.0)^42^ was utilized to characterize potential druggable binding sites. The druggable pocket score cutoff was set to 0.5. Because no druggable pockets of 39 GPCRs are identified in AlphaFold2-predicted models, the druggable pockets of only 369 AlphaFold2-predicted GPCRs were obtained. 625 potential pockets of AlphaFold2-predicted models were predicted. The number of predicted pockets in PDB structures and homology models are 764, 1867, respectively. Molecular docking was processed by AutoDock Vina (version 1.1.2)^99^. To improve the searching space, the exhaustiveness parameter was increased to 30. For each docking process, top 10 best binding affinities were kept in our results and only the top one-ranked scoring conformation and docking score were considered for comparison.

The top-one ranked docking scores of all ligand-GPCR pairs were extracted from docking results. For each GPCR with multiple pockets, only the pocket with the top-one ranked score was considered. In total, we achieved 1,680,096 metabolite-GPCR pairs, including 322,500 pairs from AlphaFold2-predicted models, 394,224 pairs from PDB structures and 967,372 pairs from homology models. For comparison, the docking score was normalized and fit into *0-1* by using adjusted Min-Max normalization method:

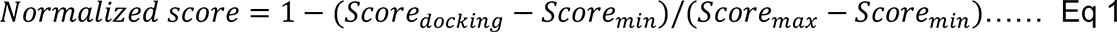

The normalization was applied to metabolite profiles to leverage the binding affinity of metabolites for each GPCR. Ward’s method was chosen as the cluster method, and Manhattan distance was adopted. The mean of normalized scores were calculated for each metabolite across all GPCRs. Heatmap was built by using R (version 2021). Detailed Binding modes and ligand-GPCR interactome were conducted by using PyMOL (version 1.8.2).

Top 10% metabolites of each GPCR were selected based on normalized score and defined them as potential metabolites. For each metabolite type and GPCR class, the percentage of potential metabolites were calculated based on the following equation:

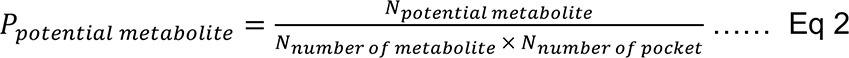

Where *P_potential metabolite_* means percentage of potential metabolites for each metabolite type and GPCR class, *N_potential metabolite_* means the number of potential metabolites for each metabolite type and GPCR class, *N_number of metabolite_* means the number of metabolites for each metabolite type, *N_number of pocket_* means the number of druggable pocket for each GPCR class.

### Machine learning framework

To improve our machine learning predict accuracy, we first calculated the physiochemical properties of metabolite datasets and took them as a reference to keep the familiar physiochemical distributions in our training dataset. Metabolites were assembled from the SMPDB database^97^ as these metabolites are associated with metabolic pathways. Molecules were cleaned by several processes, including removing ions, metals, salts, non-parsed KEGG id or names as well as super-cyclic compounds or peptides. Chemical name was converted to SMILES by using opsin (version 2.6.0). KEGG_id was converted to INCHIKEY by website (http://cts.fiehnlab.ucdavis.edu/batch). INCHIKEY was converted to SMILES by using RDKit (version 2021.03.5). Physiochemical properties were calculated by DataWarrior software (version 5.2.1)^100^, including molecular weight, hydrogen bond acceptor (HBA), hydrogen bond donor (HBD), cLogP, total polar surface area (TPSA) and Number of rotate bond (NRotB). Herein, the threshold of these six molecular properties was set to 90%, which was used to narrow down the training dataset.

Next, we assembled the bioactive ligands targeting the entire GPCR family with Ki values from GPCRdb (1,045,681 entries) and GLASS databases (321,881 entries). Duplicates, abnormal and outlier (Maximum/Minimum > 10) entries were removed. Here, we only considered activity data from human species and only bioactive ligand pairs were kept (pKi >= 6). To mimic properties of metabolites, the metabolites dataset cutoff was applied, leading to a total 60,356 ligand-GPCR entries in our training set. Molecular docking was conducted for all entries, and the detailed docking methods are described in the Molecular docking and interaction analysis section.

Then, we calculated 3D interaction features based on our docking results. All features of each ligand-GPCR pair employed in our development of the ML model was calculated by using deltaVinaXGB (https://github.com/jenniening/deltaVinaXGB) as described in a previous study^81^. Notably, because the training dataset utilized unique ligand-GPCR pairs, we removed ligand dependent features. Finally, 84 features were calculated, including 52 features describing ligand-interaction terms calculated from Vina source code, 29 features describing buried solvent accessible surface area (bSASA) and pharmacophore-based bSASA terms, 2 features describing ligand conformation stability (ligand RMSD and ΔE) during the binding process and 1 docking score. Docking score converted to pKvina following this formula: pKvina = −0.7335 ∗ Docking Score. Finally, all features of 515 metabolites were calculated.

Due to data loss upon molecular docking and feature calculation, a total of 54,446 pairs with docking score and all features, were used to construct our dataset for ML models. 10% dataset was chosen as validation dataset, the rest is training dataset. ML models were built by using PyCaret library^101^. p*Ki* value is our predicted label by using regression. The dataset was randomly divided into training set and test set with a ratio 7:3, and they were preprocessed to remove multicollinearity features with a threshold of 0.95. 10-Fold cross-validation was adopted. ML models were generated by using default hyper-parameters implemented in PyCaret. 11 regression ML models were predicted as listed on **Supplementary Table 1**^101^, including Bayesian Ridge, CatBoost, Elastic Net, Extreme Gradient Boosting, Extra Trees, Linear Regression, Light Gradient Boosting Machine, Gradient Boosting, Random Forest, Ridge Regression, K neighbors. Specifically, for Extra Trees model, random seed = 223, n_estimators = 100, min_samples_split = 2. To comparatively evaluate the ML models, Pearson’s correlation coefficient (*R*) between experimental measured binding affinity (p*Ki*) and predicted affinity was adopted to assess their linear correlation in regression. The final model was used to predict our metabolite-GPCR pairs dataset and generate the predicted GPCR-ML score (**Supplementary Data 3**).

### Benchmark study

Datasets of reported metabolite-GPCR for the benchmark study were collected from GPCRdb and high-throughput functional screening report^7^. To keep a high confidence of the dataset, only agonists determined from a β-Arrestin assay were considered. Further, agonists with pEC50 that are less than 6 were removed. Finally, 56 metabolite-GPCR pairs that comprised of 34 GPCRs and 44 metabolites were processed by structure preparation, molecular docking, feature calculation and ML model prediction.

### Mendelian Randomization analysis

We performed Mendelian Randomization (MR)^102^ to test the causal effect of 409 GPCRs toward AD. Publicly available 4 cis-eQTL (Expression quantitative trait locus) datasets of brain cortex region were used as instrument variables source in MR analysis. They were downloaded from AD Knowledge Portal (Synapse ID: syn17015233)^103^ and MetaBrain^104^. We used two criteria to select high confidential instrumental variables for each GPCR: (1) FDR < 0.05 as the cutoff of effect instrumental variables from eQTL datasets; (2) we clumped LD (linkage disequilibrium) to r^2^ < 0.2 based on reference matrices from 1000 genome V3 by PLINK [1.9]^105^. We used 3 publicly available AD GWAS summary statistic datasets as AD outcomes, including 2 GWAS from late onset AD^30, 31^ and a GWAS from the cohort with family history of AD^29^. The MR method Wald ratio estimator^106^ was used with one variant, while the inverse-variance-weighted (IVW)^107, 108^ method was used to test the number of proposed variables. The MR analysis was conducted via TwoSampleMR (https://mrcieu.github.io/TwoSampleMR/) and Mendelian Randomization packages^109^ using the R (R 4.1.1) platform. The analyzed data is available in **Supplementary Data 4**.

### Multi-omics analysis

We used 88 bulk and single cell RNA-seq transcriptome or proteome datasets that we previously compiled^35^ (available from AlzGPS: https://alzgps.lerner.ccf.org/) and examined the differential expression of the GPCRs in these datasets. Each dataset compares a pathology group (e.g., AD) and control group (e.g., healthy control). Differential expression in one or more datasets was considered evidence for the GPCRs. The detailed method could be referenced our previous study^35^. The analyzed data is available in **Supplementary Data 5**.

#### Differential expression analyses from brain transcriptome analyses

The transcriptome analyses were performed based on microarray, bulk RNA-seq, and single-cell/nucleus (sc/sn) RNA-seq datasets. We utilized three sets of human brain microarray transcriptome data collected from late-stage AD and control donors. The original data are available from Gene Expression Omnibus database (Edgar et al., 2002): (1) GSE29378 with 31 late-stage AD and 32 controls, (2) GSE48350 with 42 late-stage AD and 173 controls, and (3) GSE84422 with 328 late-stage AD and 214 controls. All results of differential expression analyses are available from the AlzGPS database^110^. We also included human brain bulk RNA-seq transcriptome data collected from hippocampus region of late-stage AD and control donors with three studies: (1) 4 late-stage AD versus 4 controls^111^, (2) 6 late-stage AD versus 6 controls^112^, and (3) 20 late-stage AD versus 10 controls^113^. For all differentially expressed genes (DEGs) generated from sc/sn RNA-seq datasets, we applied uniform criterion with adjusted p-value (q) < 0.05 and |log_2_FC| ≥ 0.25. Finally, for the sake of convenience, we have included the complete lists of DEGs based on microarray and bulk RNA seq in **Supplemental Data 5**.

#### Differential expression analyses from brain proteomic analyses

Differential expression analyses of brain proteomic data were conducted based on six mouse model datasets: (1) 7- and 10-months ADLP mouse models (JNPL3 mouse model cross with 5xFAD mouse model)^114^, (2) 7- and 10-months 5xFAD mouse models^114^, (3) 12-months 5xFAD mouse model^115^, and (4) 12 months hAPP mouse model^115^. The differentially expressed proteins (DEPs) for across different mouse brain are available from AlzGPS^110^. The complete lists of DEPs are provided in **Supplemental Data 5**.

#### Differential expression analyses of brain single-cell/nucleus RNA-sequencing data

We utilized one set of human brain single nucleus RNA-sequencing data collected from 18 AD and control donors with two brain regions: occipital cortex (OC) and occipitotemporal cortex (OTC) which included totally 482,472 nuclei. It is available from Gene Expression Omnibus (https://www.ncbi.nlm.nih.gov/geo/) database with accession number GSE148822. We performed the bioinformatics analyses according to the processes described in the original manuscript^55^. We first used DoubletFinder^116^ to remove doublets for each individual samples. After that, the rest analyses were implemented with Seurat (4.0.6)^117^, nuclei with ≤ 500 and ≥ 2500 genes, and with ≥ 5% mitochondrial genes were removed. Then the raw count was log-normalized and the top 2000 most variable genes were detected by function *FindVariableFeatures* with *selection.method* = ‘*vst’*. Next, all samples were integrated by functions *FindIntegrationAnchors* using reciprocal PCA (RPCA) and *InegrateData* with default parameters. We then scaled the data and regressed out heterogeneity related with mitochondrial content, sex, and number of UMIs. Principal component analysis (PCA) was performed with parameter *npcs = 40*, and clustering was performed with the first 13 pcs and resolution 0.15. After identifying microglia nuclei with marker genes (*CSF1R*, *C3, CIITA, P2RY12* and *CX3CR1*) provided by the original manuscript^118^, we continued with the microglia subcluster analysis with top 1000 most highly variable genes detected by function *FindVariableFeatures*. All microglia nuclei were integrated again via functions *FindIntegrationAnchors* using canonical correlation analysis (CCA) with parameters *dims = 1:10*, and *IntegrateData* with default parameters. After that, data was scaled and regressed out ribosomal content. We then ran the subcluster partition with the first 10 pcs and resolution 1.3. We identified different microglia subtypes with the markers provided in the original manuscript^118^: *TMEM163*, *CX3CR1*, *SOX5*, and *P2RY12* for homeostasis microglia, *ITGAX*, *SPP1*, *MSR1*, and *MYO1E* for Aβ related microglia, *GRID2*, *ADGRB3*, *CX3CR1*, and *DPP10* for Tau related microglia, *GPNMB*, *IL1B*, *CD83* and *NFKB1* for inflammation microglia, *HSP1A1*, *HSPA1B*, *FOS*, and *HSP90AA1* for stress microglia, and *TOP2A*, *BRIP1*, *MKI67*, and *FANCI* for proliferation microglia. DEGs were calculated between AD1, AD2, inflammation and homeostasis microglia with MAST R package^119^, separately. The analyzed data is available in **Supplementary Data 6**.

### Interactome network modeling

The Metabolite-GPCR interaction network was visualized using Cytoscape (version 3.9.1). To build this network, we used several layers of our results: (1) 884 nodes, including 515 metabolites and 369 GPCRs with their classification; (2) 884 edges, includes 515 metabolite-GPCR pairs between metabolites with strongest GPCR-ML score and their targeted GPCR, and 369 metabolite-GPCR pairs between GPCRs and their top-one ranked metabolites; (3) MR and multi-omics evidence of GPCRs in AD; (4) Microbiota bacteria strains related to each metabolite. Metabolites were classified by chemical types; GPCRs were classified by ligand types (**Supplementary Data 7**). The metabolic profiles across bacteria strains were from re-analysis of previous study^40^ (**Supplementary Data 9**).

### Pocket classification and analysis

Since different GPCR classes have their unique pocket characteristics, we need to recognize the orthosteric or allosteric pocket for each GPCR class. To define the relative position of pocket, the generic residue number was adopted based on GPCRdb numbering scheme^41^.

For class A, the traditional orthosteric site was considered as being located in extracellular sites between the transmembrane helix segments. As in previous report^120^, several relatively high frequency residues are regarded as our reference residues: 2×60, 2×63, 3×28, 3×29, 3×32, 3×33, 3×36, 3×37, 45×52, 5×40, 5×41, 5×44, 5×45, 5×47, 6×44, 6×48, 6×51, 6×52, 6×55, 6×58, 7×30, 7×33, 7×37, 7×40, 7×41. Classified the pocket as an orthosteric pocket once the binding pocket possesses at least 5 residues among the reference residues. At first, generic numbers of reference residues are matched to sequence numbers of each protein model. Then we applied the above condition to each pocket in each model, leading to a set of pocket types. Next, we compared every docking pocket of GPCR-metabolite pairs with the pocket set, and finally we could analyze the orthosteric pocket distributions in docking results. A pocket that is different from identified orthosteric pocket was regarded as an allosteric pocket. The same analysis was employed to class T receptors^121^.

For class B receptors, since they are peptide receptors, the orthosteric pocket would be much larger and more solvent accessible^122^. Therefore 36 complex structures with ligand binding to orthosteric sites of class B were investigated and residues involved in interactions are analyzed. Then we selected the residues with frequency of more than 50%, and finally 26 residues are regarded as the pocket reference residues. Next, we match the reference residues to each pocket and compare them in docking datasets. Same analysis was employed for class F receptors^123^.

Class C receptors differ from other subfamilies. They are structurally distinguished by a large N-terminal extracellular domain, and endogenous ligands are identified by binding this domain^124^. Since it is difficult to obtain the generic number of N-terminal residues, we inspected all pocket positions and differentiated among them manually. We then utilized them and distinguished the pockets in docking datasets.

In total, 625 pockets from AlphaFold2, 767 pockets from PDB and 1867 pockets from GPCRdb were analyzed and applied to define pocket types on all GPCR-metabolites docking pairs. (**Supplementary Data 8**)

### GPCRome Phylogenetic tree

Sequences of 408 GPCRs were collected from UniProt (https://www.uniprot.org/). The multiple alignment sequence (MSA) information was achieved from GPCRdb. Only structurally conserved positions and TM1-TM7 segments were considered to construct the alignment file. Finally, the MSA with 393 GPCRs was used to construct the phylogenetic tree by utilizing Phylip (version 3.695). Furthermore, the number of transcriptome and proteome evidence were plotted on the outer circle. The circular tree was polished by using iTOL (version 6.5.2).

### Statistical analysis

All statistical analyses were performed using the R package 4.1.1.

### Data availability

Genome-wide association studies (GWAS), transcriptomics, and proteomics datasets were collected from The National Institute on Aging Genetics of Alzheimer’s Disease Data Storage Site (NIAGADS, www.niagads.org) and The Alzheimer’s disease Knowledge portal (https://adknowledgeportal.synapse.org/). The data that support the findings of this study are available in the manuscript and its supplementary information. The list of studied metabolites and GPCRs are provided in **Supplementary Data 1-2.** The GPCR-ML scores of metabolites on AlphaFold2-predicted models, homology models and PDB models are provided in **Supplementary Data 3**. The target analysis of Mendelian Randomization (**Supplementary Data 4**), multi-omics (**Supplementary Data 5**) and Single nucleus datasets (**Supplementary Data 6**) are available in **Supplementary Data 4-6**. The interactome associations of metabolites and GPCRs shown in Fig. 3c are available in **Supplementary Data 7**. The pocket distributions of AlphaFold2-predicted models, homology models and PDB models are available in **Supplementary Data 8**. The re-analysis of profile of metabolites across bacterial strains are available in **Supplementary Data 9**. All data are available from the authors upon reasonable request.

### Code availability

Codes for machine learning frameworks and other data analyses are available at https://github.com/ChengF-Lab/Gut-GPCRome. All other codes used in this study are available from the corresponding author upon reasonable request.

## Acknowledgments

We thank members from The National Institute on Aging Genetics of Alzheimer’s Disease Data Storage Site and The Alzheimer’s disease Knowledge portal to provide technical support for multi-omics data access and analysis.

## Author contributions

F.C. conceived the study. Y.Q., Y.H., Y.Z., J.X. performed all experiments and data analysis. M.B., J.B.L., A.A.P., R.N., J.M.B. discussed and interpreted the results. Y.Q., A.A.P., and F.C. wrote the manuscripts. All authors critically revised and gave final approval of the manuscript.

