## Supplementary Figures for "Comprehensive characterization of multi-omic landscapes between gut-microbiota metabolites and the G-protein-coupled receptors in Alzheimer’s disease"

**Contents:**

**Supplementary Figures 1-17, Table 1-3 and legends.**

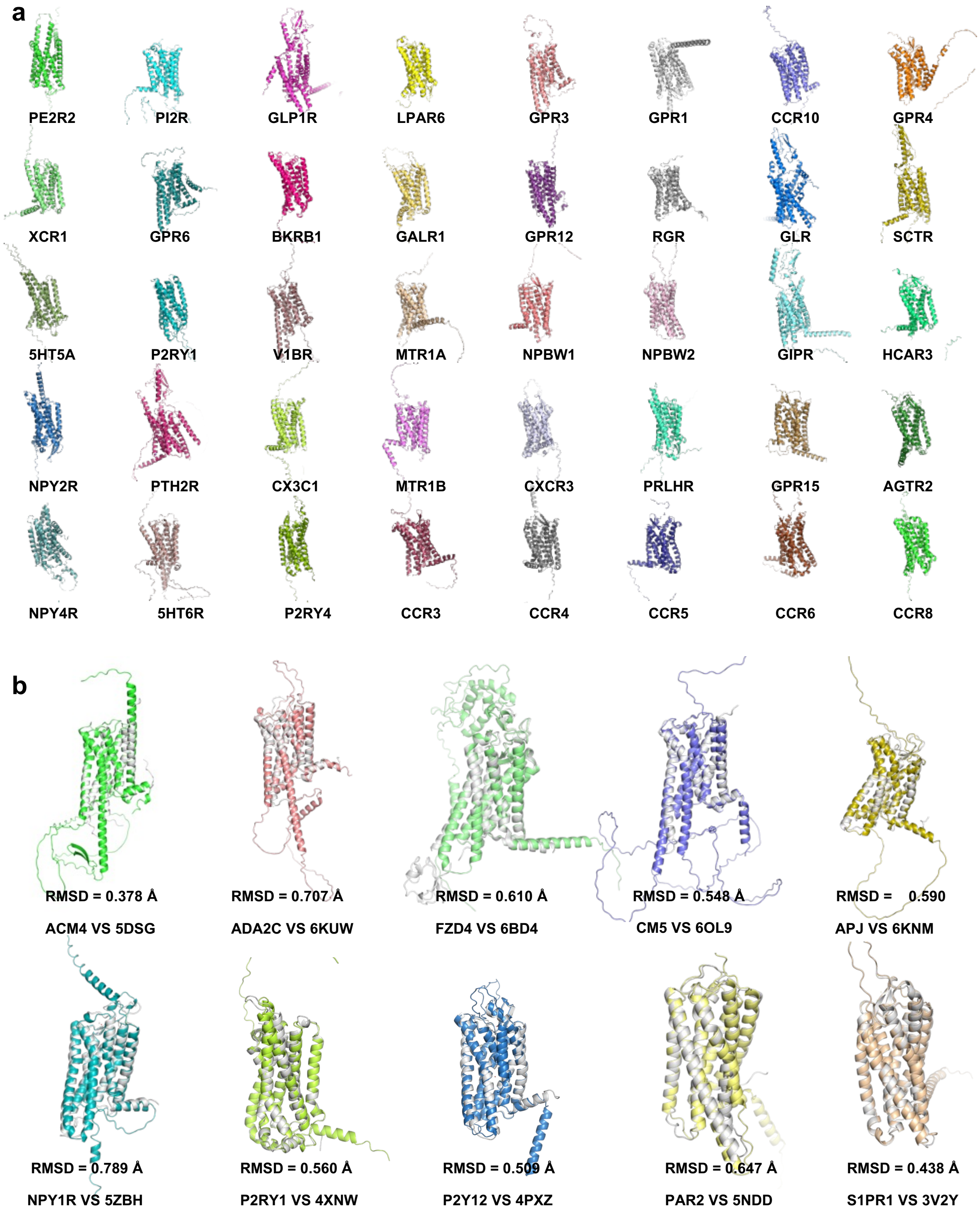


**Supplementary Figure 1. Diagram showing 3D structures of 40 AlphaFold2-predicted GPCR models and comparisons of 10 AlphaFold2-predictied models with crystal structures.** The GPCRs are random selected from GPCRome models. RMSD between AlphaFold2 and crystal structures are displayed, AlphaFold2 models were shown in different color, while crystal structures shown in gray cartoon.

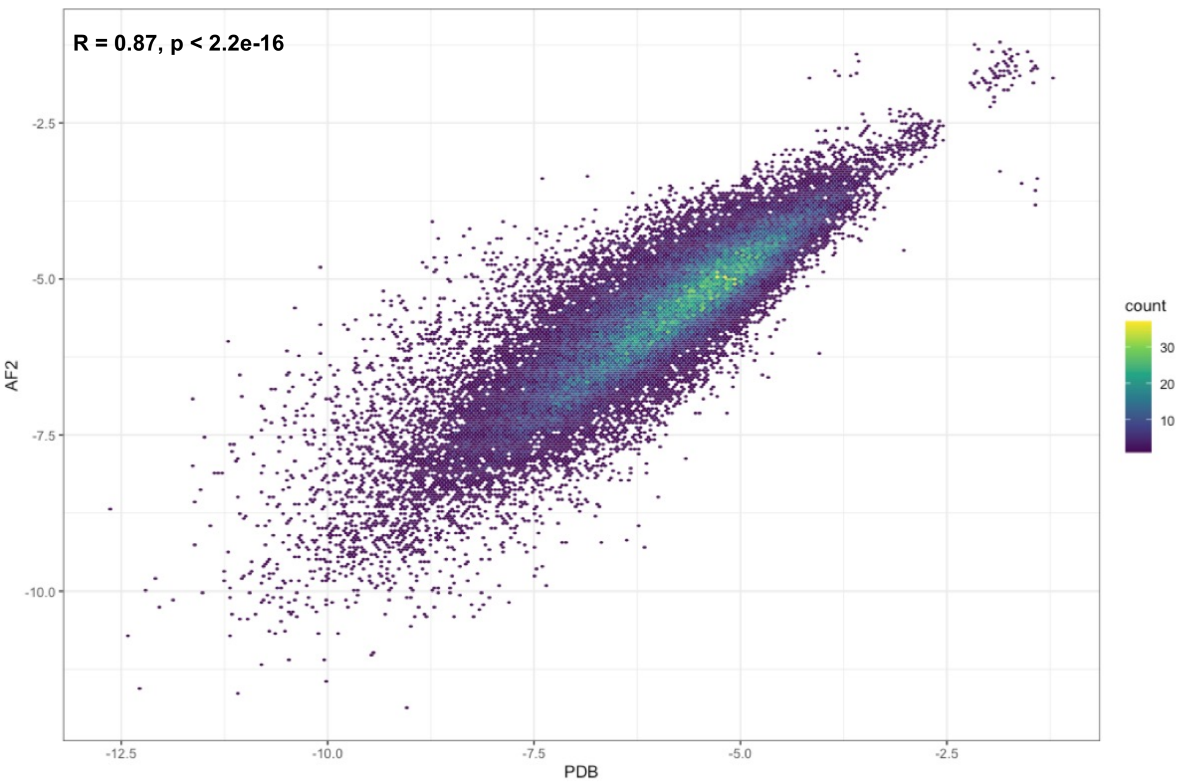


**Supplementary Figure 2. Docking score correlation between 77 identical GPCRs of PDB structures and AlphaFold2-predicted models.** Metabolite-GPCR pairs were scatter plotted with density in different color. Pearson's correlation coefficient R and *p* value were labeled.

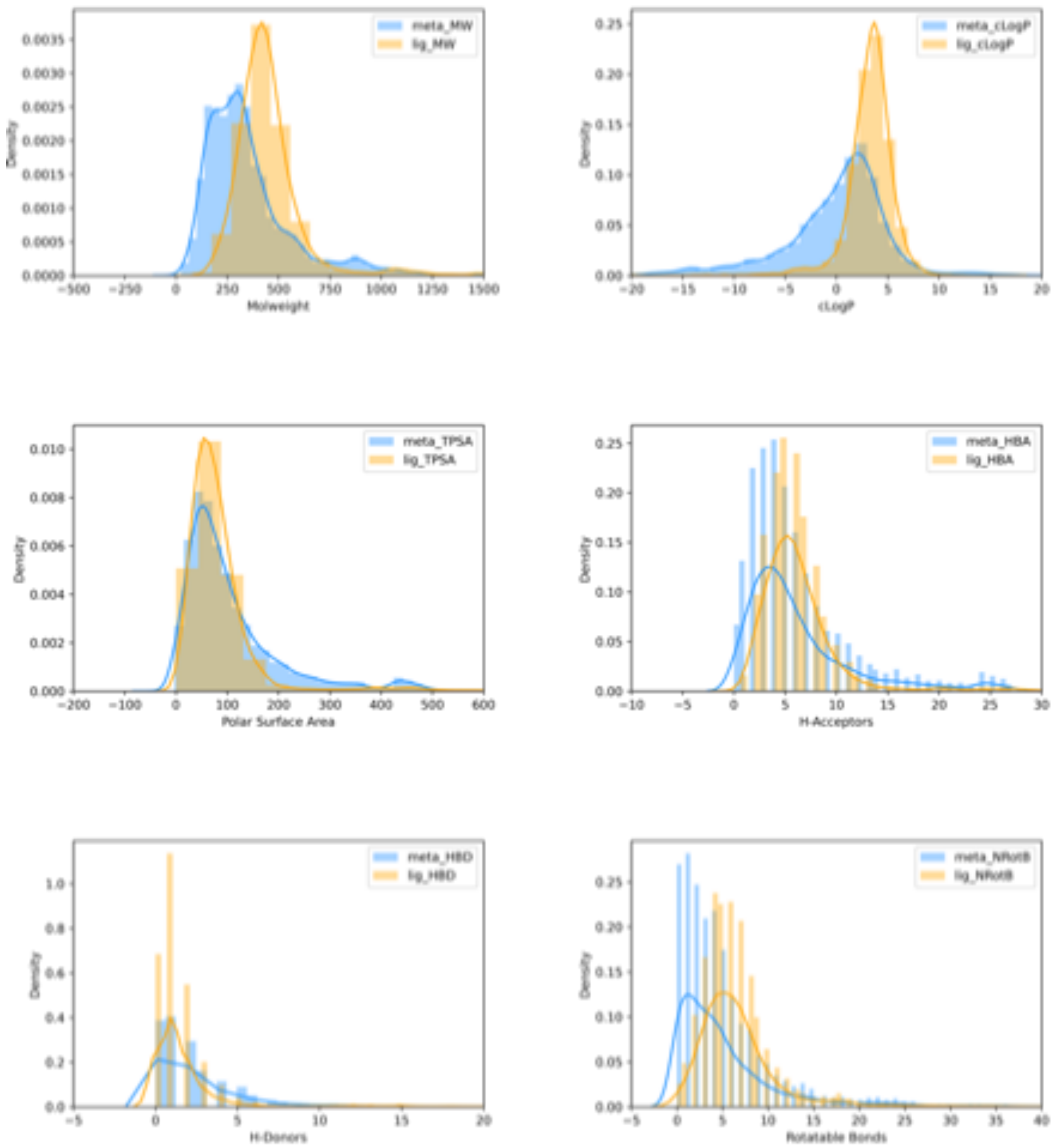


**Supplementary Figure 3. Frequency analysis of physicochemical properties.** Six physicochemical properties, including Molecular Weight, cLogP, Polar Surface Area, H-Acceptor, H-Donor and Rotatable bonds, were calculated and compared between reported GPCR bioactive compounds from GPCRdb and GLASS database and metabolites from SMPDB database. Property frequencies of bioactive compound dataset are depicted in orange, while that of datasets are depicted in blue.

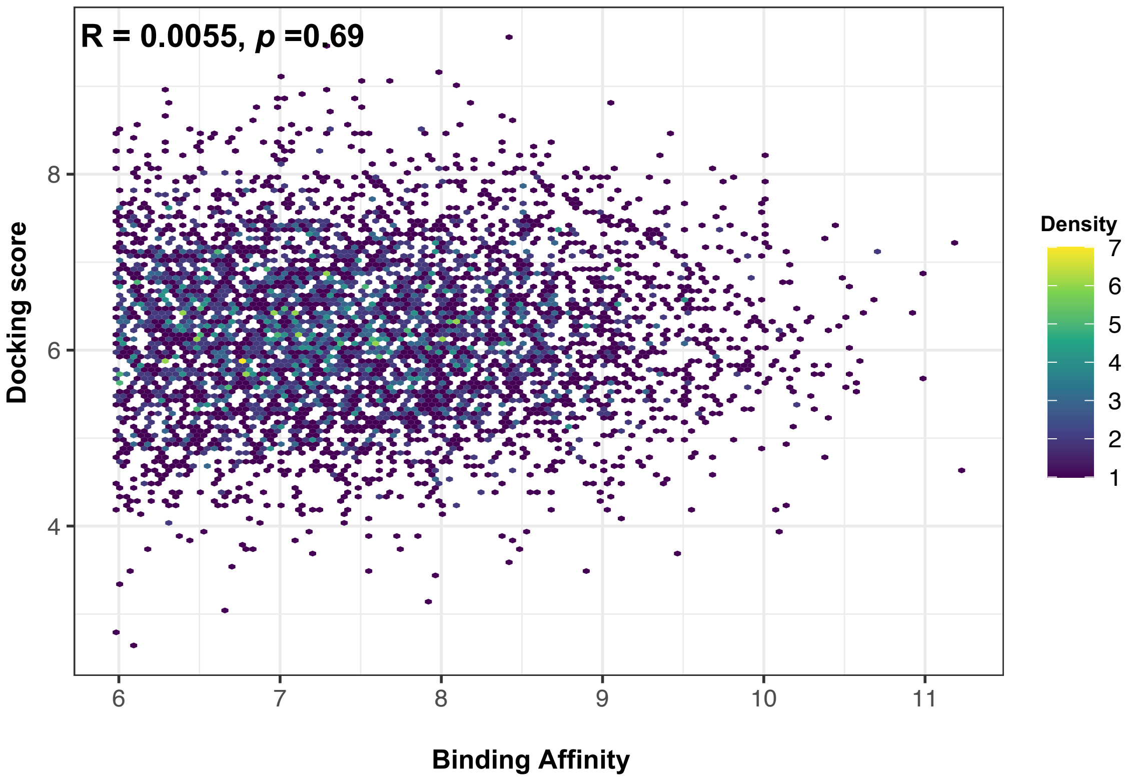


**Supplementary Figure 4. Performance of molecular docking on external test dataset.** Binding affinity and docking score of 5445 unseen metabolite-GPCR pairs were scatter plotted with density in different color. Pearson's correlation coefficient R and *p* value were calculated.

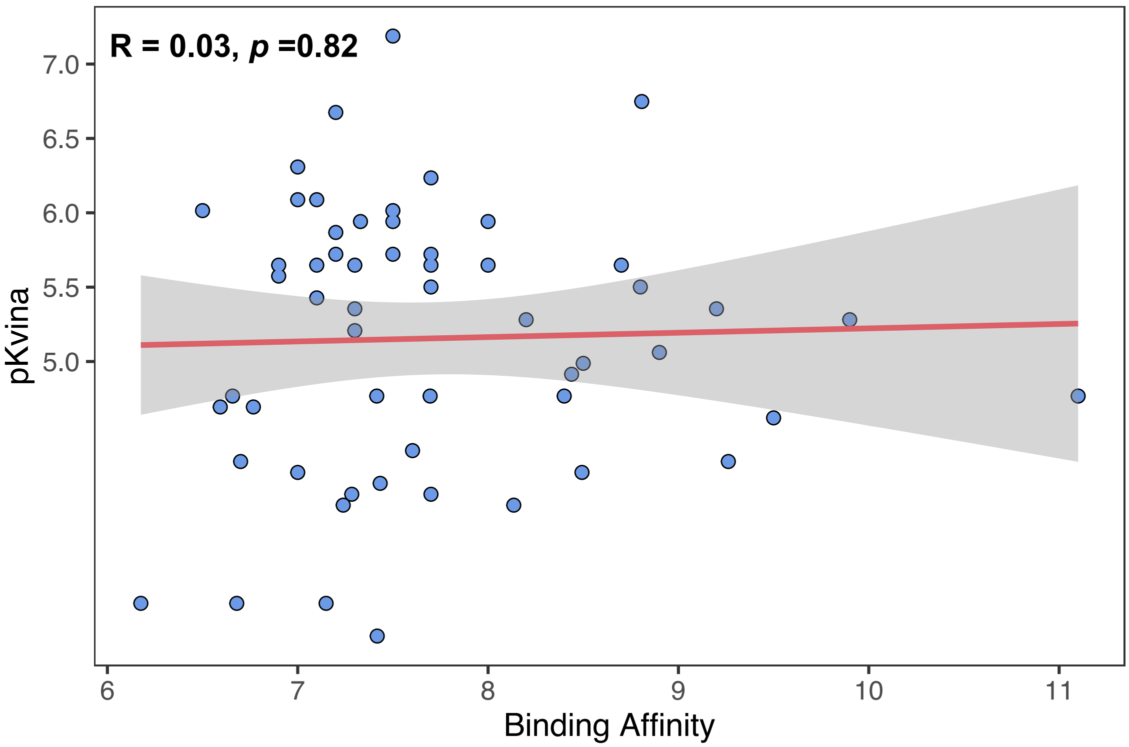


**Supplementary Figure 5. The performance of molecular docking on benchmark dataset.** Binding affinity and docking score of 56 reported metabolite-GPCR pairs were scattered plotted in color blue. Regression of data was predicted by Linear Regression function, displayed as red line along with SD error (gray background). Pearson's correlation coefficient R and *p* value were labeled.

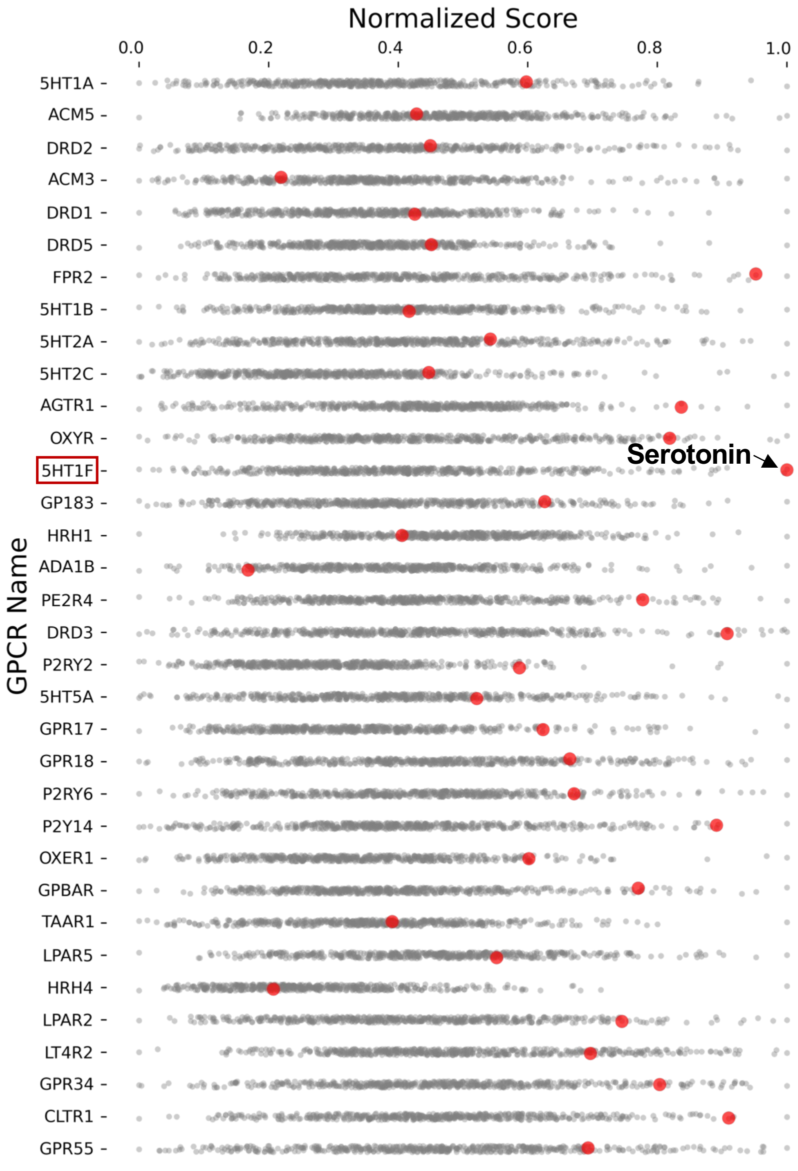


| GPCR Name | pEC50 | GPCR-ML Score | Absolute Error |
|---|---|---|---|
| 5HT1A | 7.23901993 | 7.44551857 | **-0.2064986** |
| ACM5 | 6.67970828 | 7.18659846 | **-0.5068902** |
| DRD2 | 7.43323264 | 7.12098527 | **0.31224738** |
| ACM3 | 6.17586114 | 7.04246234 | **-0.8666012** |
| DRD1 | 6.65765629 | 7.25692718 | **-0.5992709** |
| DRD5 | 6.76702929 | 7.31932708 | **-0.5522978** |
| FPR2 | 11.1 | 7.92431708 | **3.17568292** |
| 5HT1B | 7.41529571 | 7.17528928 | **0.24000644** |
| 5HT2A | 7.69559565 | 7.29394737 | **0.40164828** |
| 5HT2C | 8.43886282 | 7.35915099 | **1.07971184** |
| AGTR1 | 9.26229859 | 7.93925134 | **1.32304726** |
| OXYR | 8.80729319 | 7.72343063 | **1.08386256** |
| 5HT1F | 7.60312097 | 7.88943656 | **-0.2863156** |
| GP183 | 7.3 | 7.53633251 | **-0.2363325** |
| HRH1 | 7.14902924 | 7.15629506 | **-0.0072658** |
| ADA1B | 7.2839382 | 6.76627828 | **0.51765992** |
| PE2R4 | 8.8 | 7.81155637 | **0.98844363** |
| DRD3 | 8.13500861 | 7.73131659 | **0.40369203** |
| P2RY2 | 7.1 | 7.74275358 | **-0.6427536** |
| 5HT5A | 8.4934138 | 7.26347727 | **1.22993653** |
| GPR17 | 9.5 | 7.63559649 | **1.86440352** |
| GPR18 | 7.7 | 7.38939539 | **0.31060462** |
| P2RY6 | 6.59303425 | 7.54132016 | **-0.9482859** |
| P2Y14 | 7.2 | 7.88972723 | **-0.6897272** |
| OXER1 | 7.7 | 7.72672144 | **-0.0267214** |
| GPBAR | 7.5 | 7.67761354 | **-0.1776135** |
| TAAR1 | 7 | 7.18068711 | **-0.1806871** |
| LPAR5 | 7.3 | 7.39994435 | **-0.0999443** |
| HRH4 | 7.41769795 | 7.08190361 | **0.33579435** |
| LPAR2 | 8.2 | 7.62824808 | **0.57175192** |
| LT4R2 | 7.5 | 7.46169079 | **0.03830921** |
| GPR34 | 6.9 | 7.72021444 | **-0.8202144** |
| CLTR1 | 7.7 | 7.86359353 | **-0.1635935** |
| GPR55 | 7.3 | 7.513411 | **-0.213411** |

**Supplementary Figure 6. Benchmark validation of reported GPCR-metabolite pair on whole dataset.** Scatter plot illustrating the score position of reported metabolites on screening dataset and comparison of pEC50 and GPCR-ML score. Screening results of 34 GPCRs are depicted by row. Each metabolite is indicated in gray point, and reported metabolites are indicated in red point. GPCR-ML score is normalized by adopting Min-Max normalization. The corresponding pEC50 and GPCR-ML score are listed on the right. Absolute Error between them is calculated.

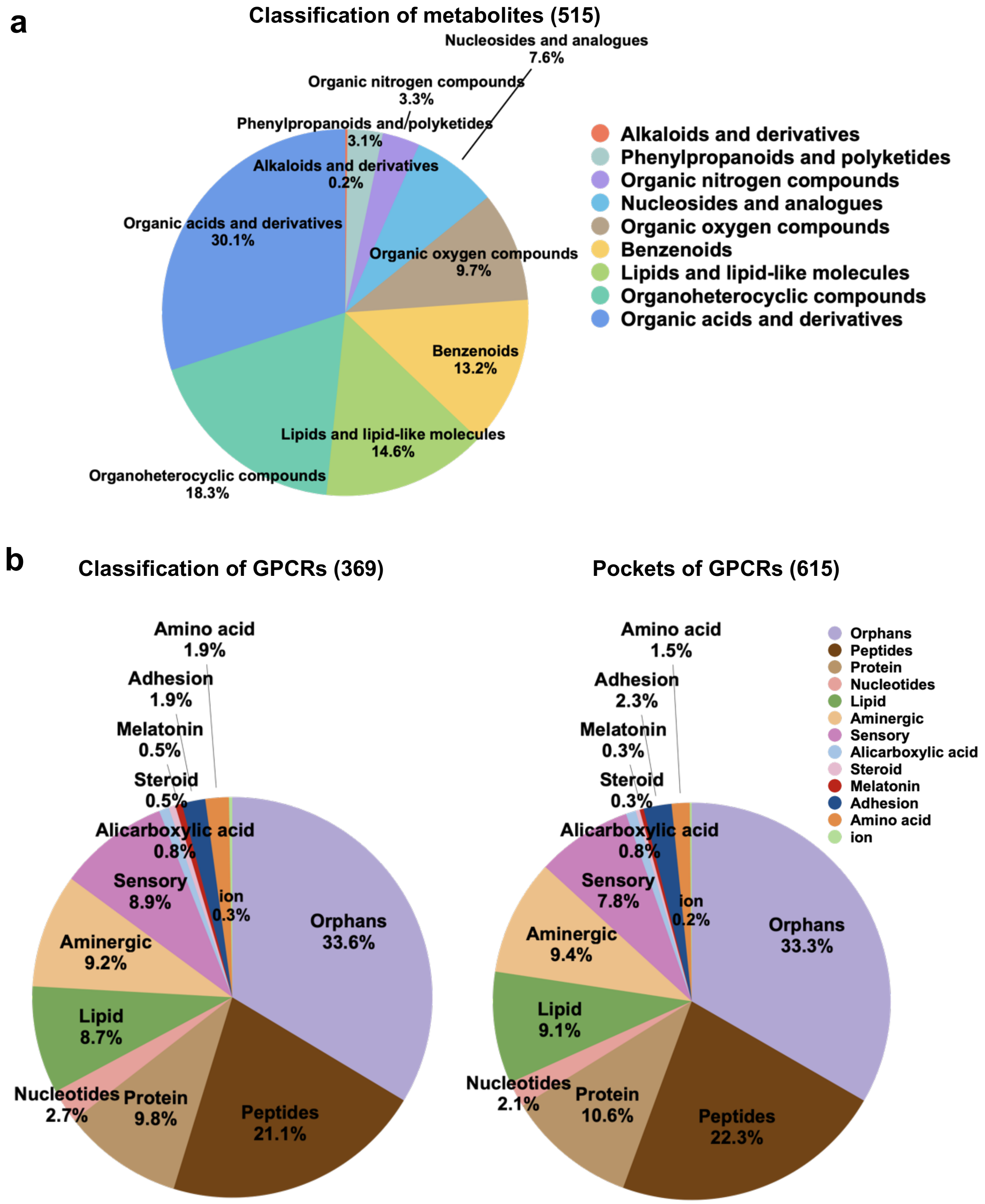


**Supplementary Figure 7. Distributions of metabolite, GPCR and GPCR pockets. a, Distributions of metabolites by chemical classes. b, Distributions of GPCRs by ligand types. c, Pocket Distributions from GPCRs by ligand types.**

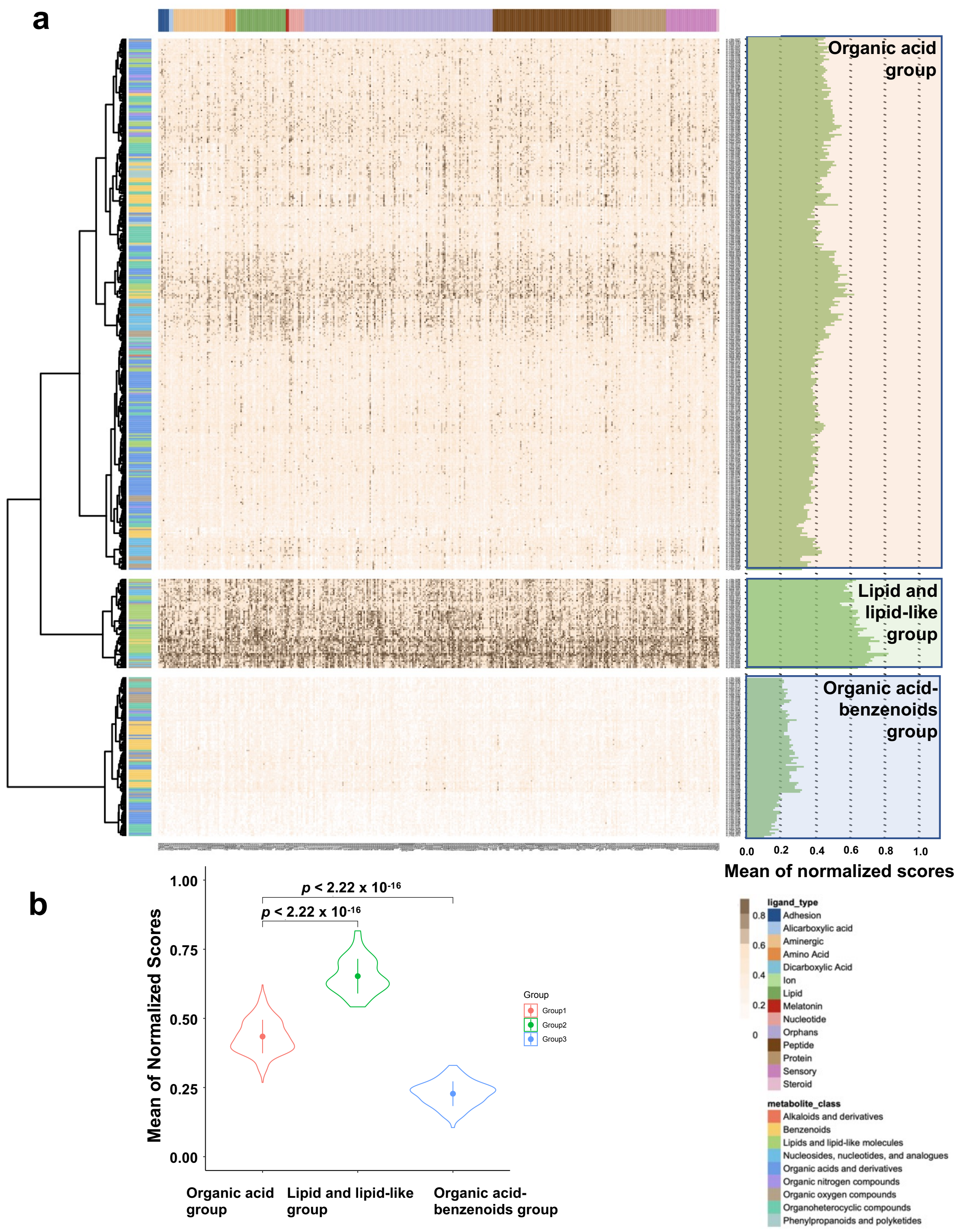


**Supplementary Figure 8. GPCR-ML score profiles of metabolite-GPCRome calculated on AlphaFold2-predicted models. a,** heatmap depicting associations between 369 GPCRs (by column) and 515 metabolites (by row). The classes of GPCR and metabolite are displayed aside with different color. GPCR-ML score is normalized by adopting Min-Max normalization. Metabolites are hierarchically clustered (Ward's D method) using Manhattan distance between the normalized GPCR-ML score across all taxonomies. The metabolite-GPCR pairs are divided into three groups based on hierarchical clusters of metabolites. For each metabolite, the mean of normalized scores across all GPCRs were calculated (by row). **b**, violin plot showing the statistical difference between organic acid group, lipid and lipid-like group, and organic acid-benzenoids group.

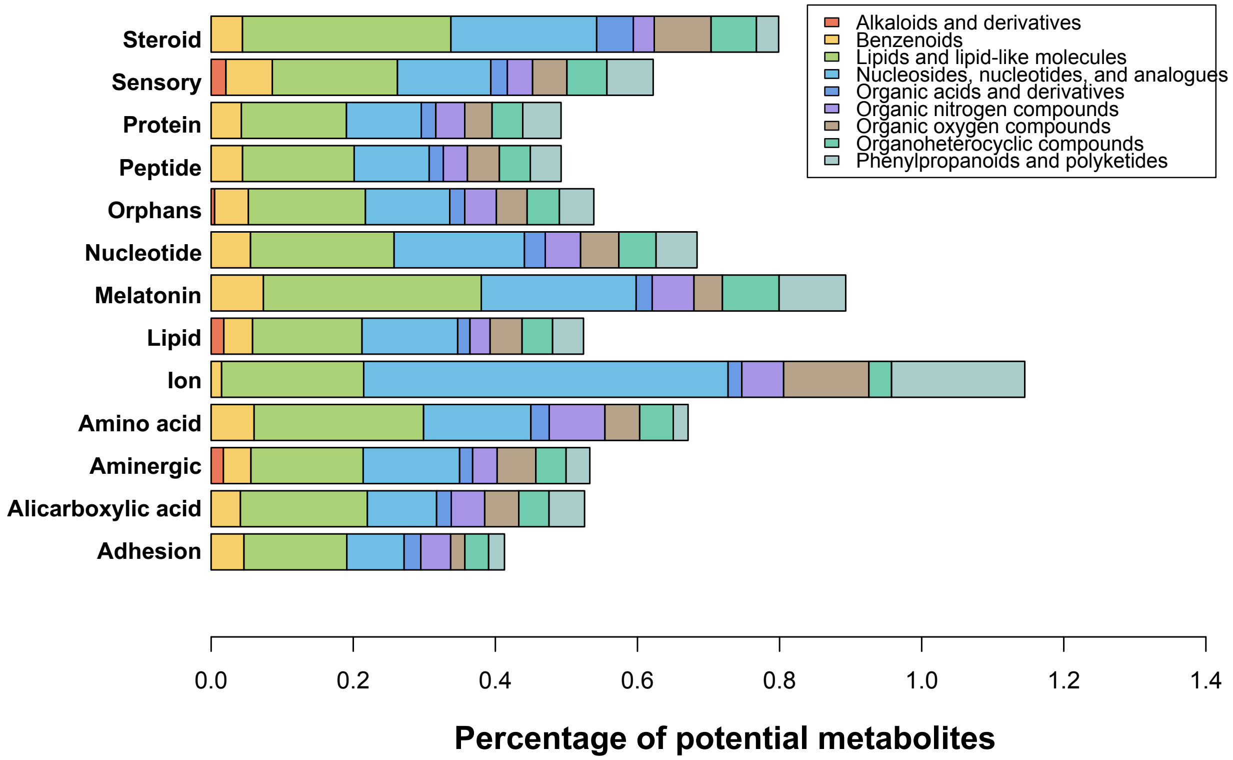


**Supplementary Figure 9. Multi-bar plot showing percentage of potential metabolites (prioritized by top 10% of metabolites ranked by normalized scores) for each type of GPCR.** For each GPCR type depicted in x-axis, the percentage of potential metabolites in different metabolite types (y-axis) was shown in multi-bar plot. Metabolite types were represented in different color.

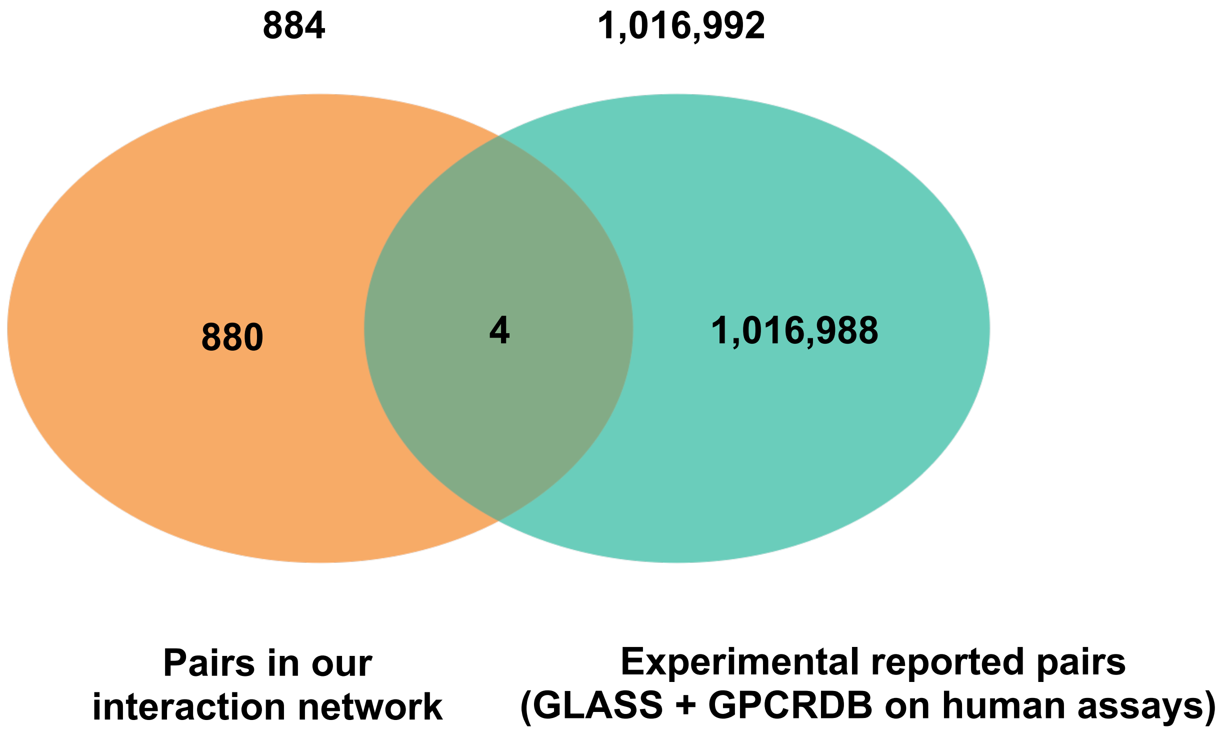


**Supplementary Figure 10. Venn diagram showing the overlap of metabolite-GPCR pairs in interactome network and reported experimental metabolite-GPCR pairs in human assays.** In total, 884 pairs were generated in our interactome network (left, orange) and 1,016,992 pairs were retrieved from GLASS and GPCRDB databases (right, green). 4 pairs were shared between two datasets.

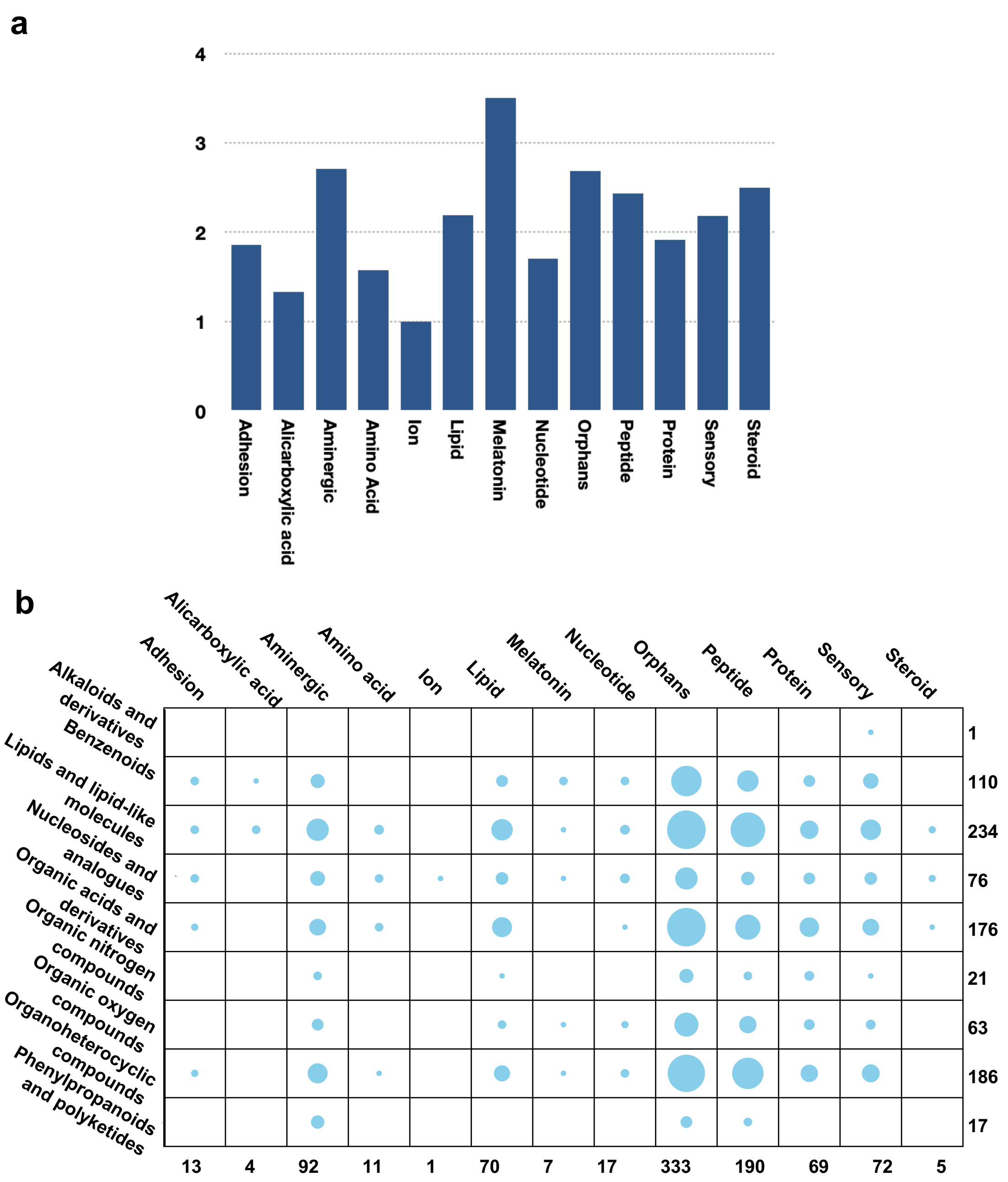


**Supplementary Figure 11. Distribution of metabolite-GPCR associations in interactome network. a,** the average number of metabolite-GPCR associations per GPCR in each GPCR class. **b,** the total number of metabolite-GPCR associations between each metabolite type and GPCR class.

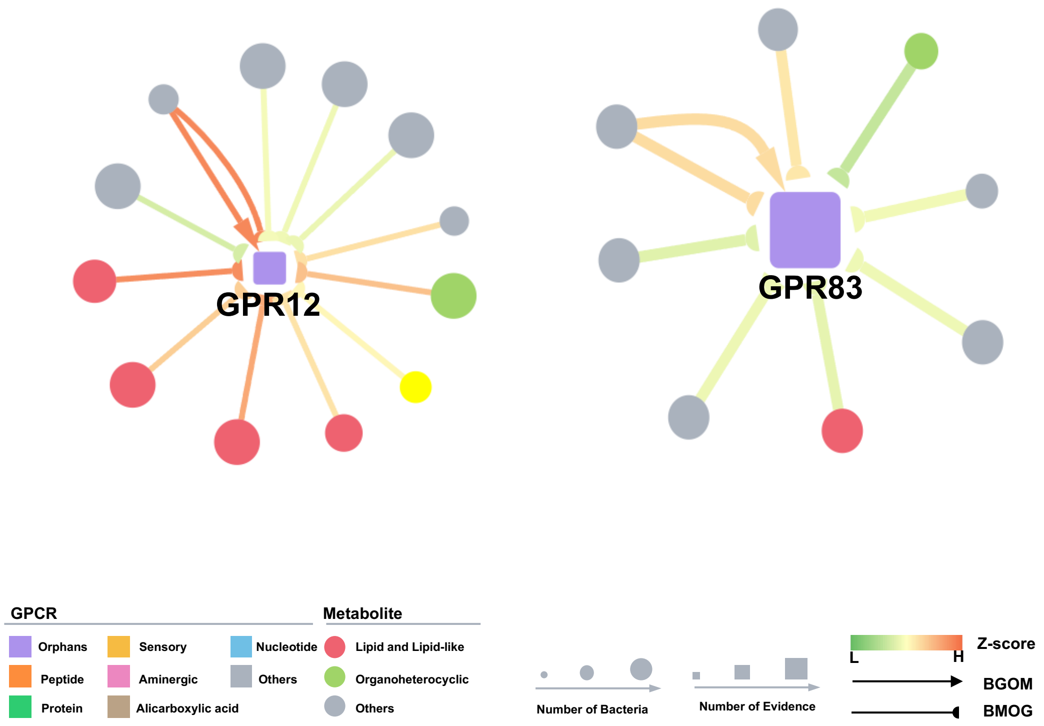


**Supplementary Figure 12. Sub-network illustrating interactions of bacteria-derived metabolites and GPR12 or GPR83.** Two sub-networks associated with GPR12 and GPR83 are displayed in detail. The top-one ranked metabolite for GPCRs or the top-one ranked GPCR for metabolites were connected by normalized GPCR-ML score. Metabolite or GPCR are separately denoted as rectangle and circle node. GPCR-ML score (edge) was normalized by z-normalization and shown in green-orange color range according to its size. The arrow edge means the top-one ranked GPCR target of metabolite, and the half-circle arrow line means the top-one ranked metabolites of GPCR. Hierarchical class of GPCR and chemical class of metabolites was indicated using different colors. The size of GPCR node is proportional to the number of MR and multi-omics evidence, while the size of metabolite node is proportional to the number of bacteria strains with higher metabolite abundance (abundance with | $Log_2FC$ | >=2 was shown).

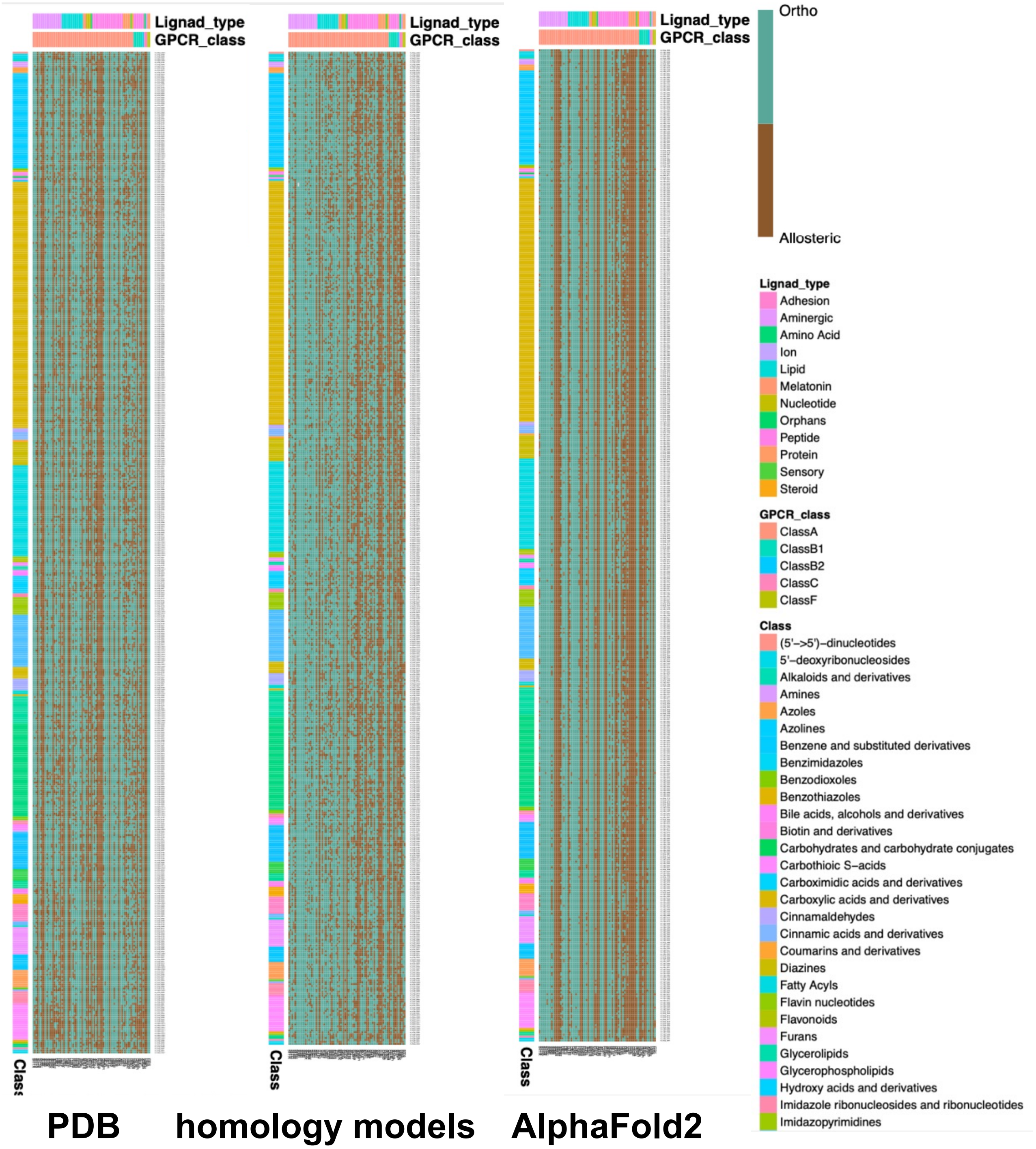


**Supplementary Figure 13. The pocket landscape of 77 identical GPCRs shared on PDB structures, homology models and AlphaFold2-predicted models.** Orthosteric pockets (green) or allosteric pockets (brown) were analyzed based on its position for all GPCR-metabolite pairs on 77 identical GPCRs shared in PDB structures, homology models and AlphaFold2-predicted models. Identical GPCRs were kept among three models and listed by column. The identical metabolites were kept among three models and listed by row. Classes of GPCR are shown in different colors based on ligand types.

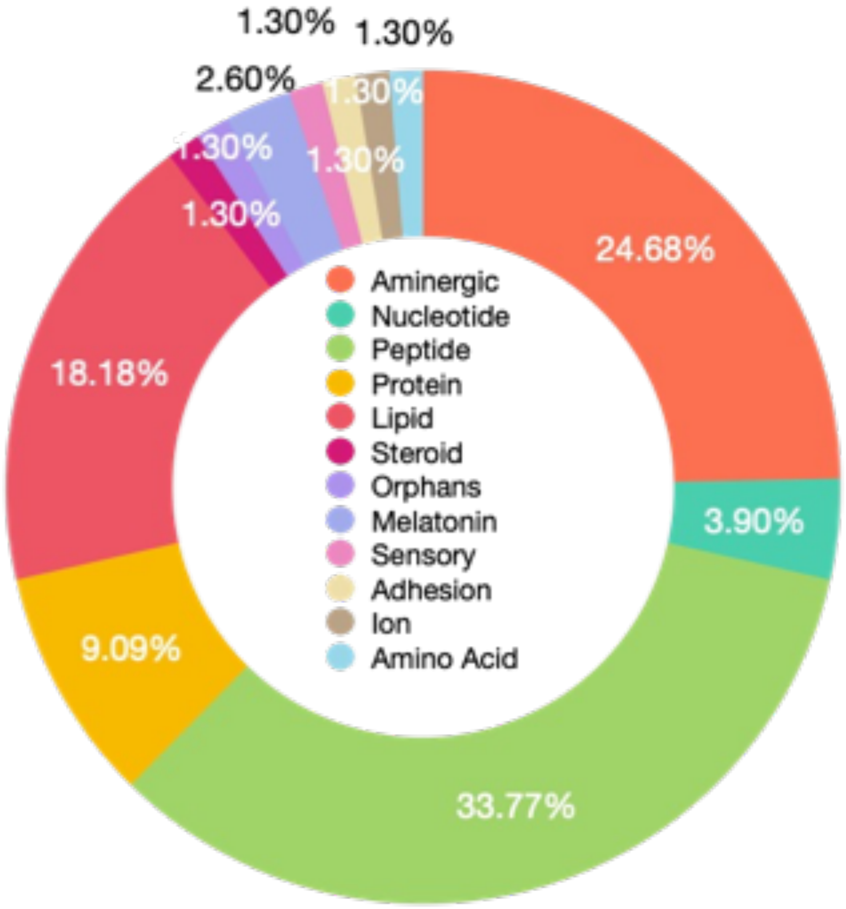


**Supplementary Figure 14. A pie chart showing percentage of GPCR class among 77 identical GPCRs shared in PDB structures, homology models and AlphaFold2-predicted models.** GPCRs were classified into 12 classes based on ligand types, and indicated by multi-color. The percentage of each class was labeled.

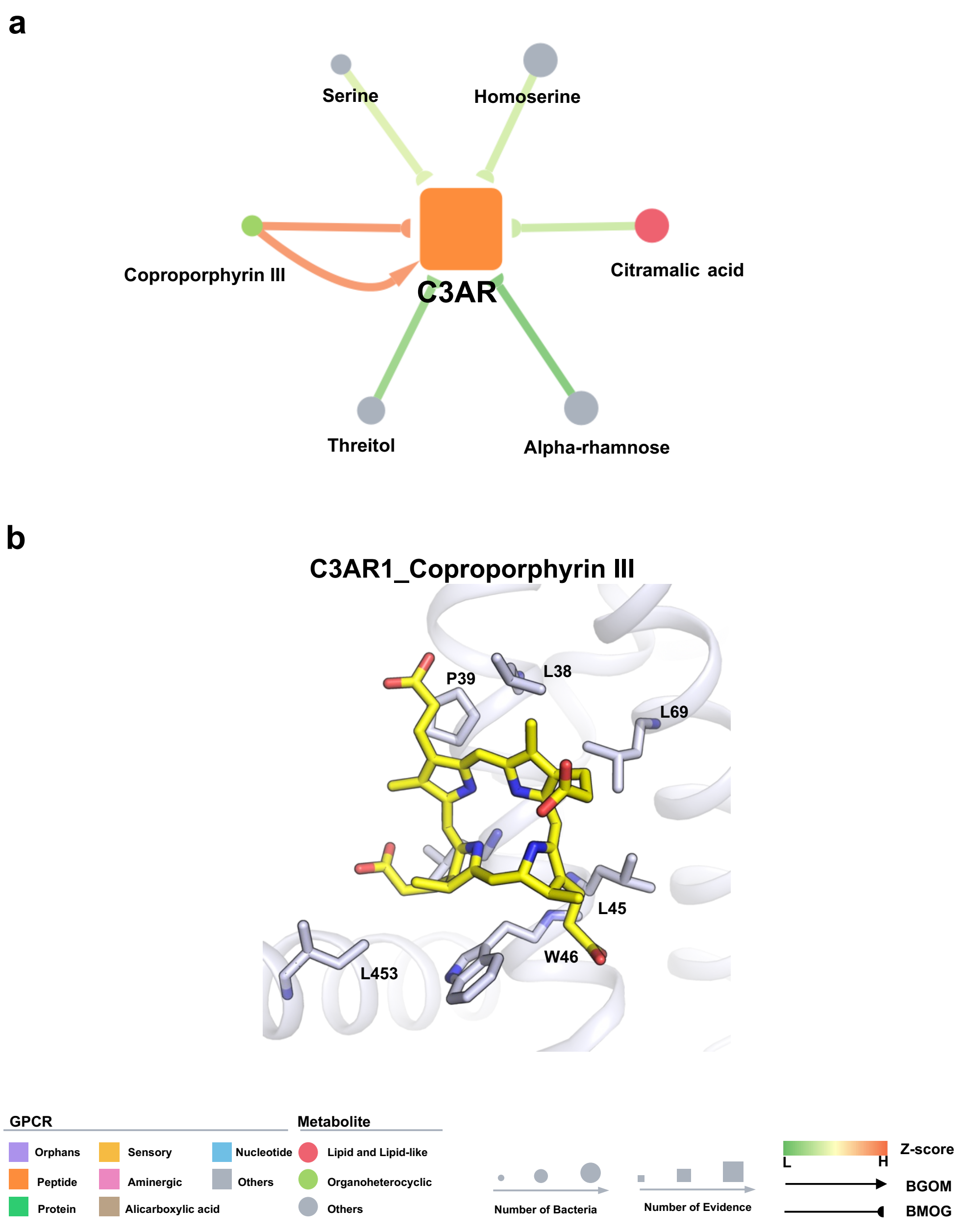


**Supplementary Figure 15. Associations of multi-omics evidenced AD-related C3AR. a**, sub-network of associations between metabolites and C3AR. The chemical names of 6 associated metabolites were labeled. The colors and lines are denoted as the same as Fig. S12. **b**, the binding mode of C3AR and its top-one ranked metabolite, Coproporphyrin III. Key residues in binding site were labeled and shown in sticks. C3AR were shown in blue cartoon.

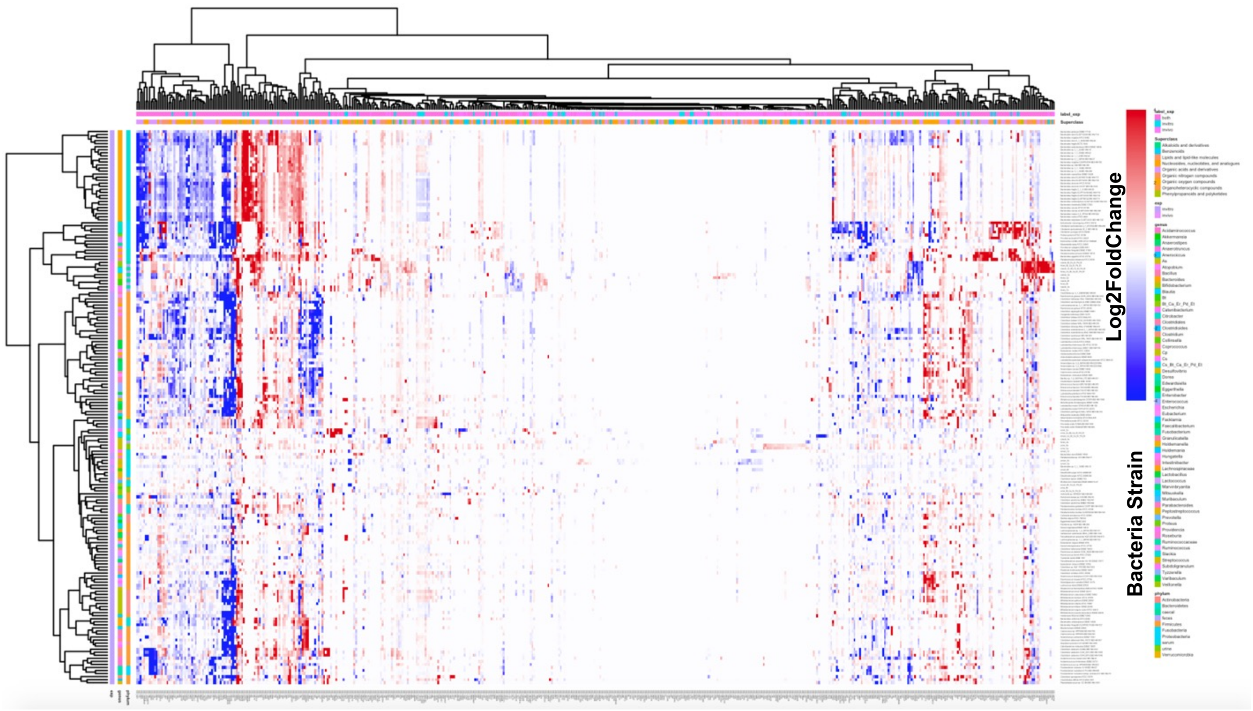


**Supplementary Figure 16. Metabolomic profiles of bacterial strains in our study. Relative Abundance of metabolites in bacteria strains compared to control was re-analyzed.** Metabolites and bacteria strains are hierarchically clustered (Ward's D method) using Manhattan distance between the Log2FC values across all taxonomies. Relative Abundance is indicated in blue-red color. For comparison, we remain the same color when | Log2FC | >=4.

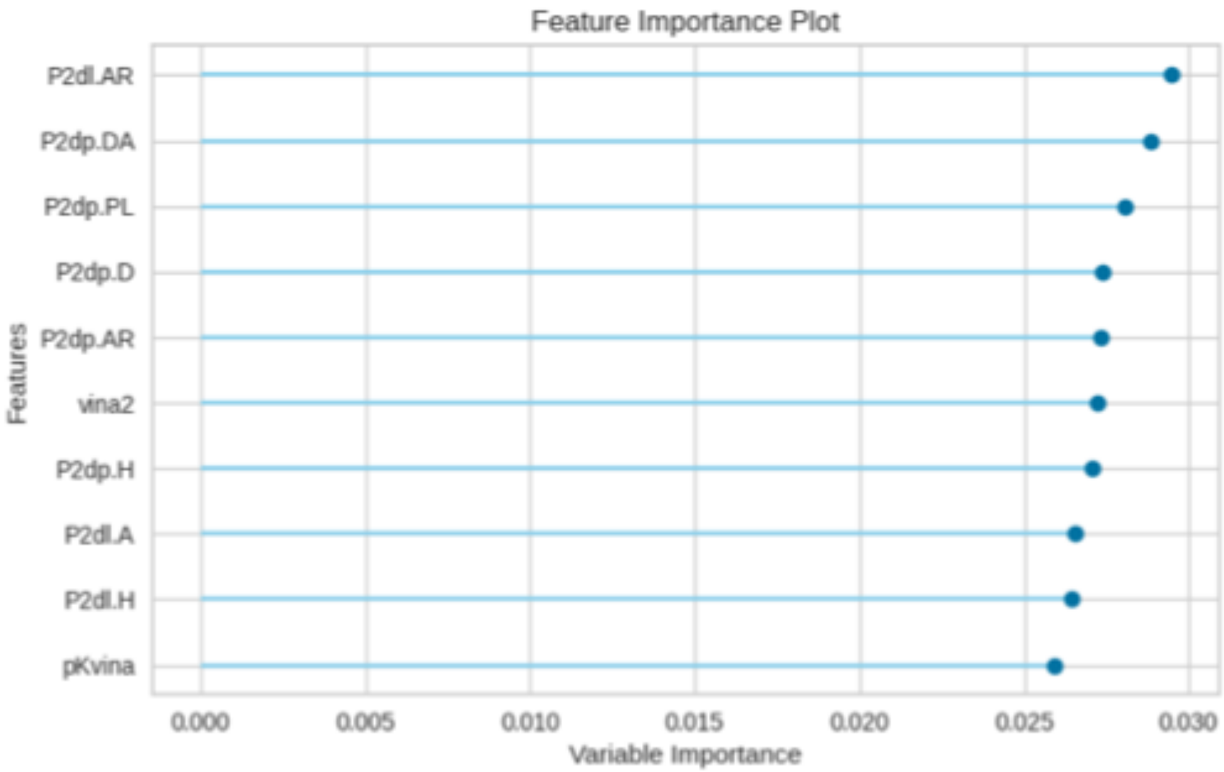


**Supplementary Figure 17. Top 10 most important features in Extra Trees model.**

**Supplementary Table 1. Prediction performance of different ML models.**

| model | MAE | MSE | RMSE | Pearson's R | RMSLE | MAPE |
|---|---|---|---|---|---|---|
| Extra Trees | 0.6308 | 0.6391 | 0.7994 | 0.603241245 | 0.0928 | 0.0847 |
| Random Forest | 0.657 | 0.6725 | 0.82 | 0.575152154 | 0.0951 | 0.0881 |
| CatBoost | 0.6918 | 0.7309 | 0.8548 | 0.522206856 | 0.0989 | 0.0924 |
| Extreme Gradient Boosting | 0.6968 | 0.7555 | 0.8691 | 0.498096376 | 0.1006 | 0.0931 |
| Light Gradient Boosting Machine | 0.7312 | 0.8 | 0.8943 | 0.451663592 | 0.1033 | 0.0976 |
| K neighbors | 0.7119 | 0.8179 | 0.9043 | 0.431161223 | 0.1048 | 0.0956 |
| Gradient Boosting | 0.7803 | 0.8979 | 0.9475 | 0.326496554 | 0.1093 | 0.104 |
| Ridge Regression | 0.8192 | 0.9822 | 0.991 | 0.150665192 | 0.1143 | 0.1091 |
| Linear Regression | 0.8193 | 0.9823 | 0.991 | 0.150332964 | 0.1143 | 0.1091 |
| Bayesian Ridge | 0.8195 | 0.9825 | 0.9911 | 0.15 | 0.1143 | 0.1092 |
| Elastic Net | 0.8223 | 0.9882 | 0.994 | 0.13 | 0.1146 | 0.1095 |

**Supplementary Table 2. 10-fold cross-validation performance of Extra Trees model.**

| C-V Fold | MAE | MSE | RMSE | Pearson's R | RMSLE | MAPE |
|---|---|---|---|---|---|---|
| 0 | 0.6314 | 0.6366 | 0.7979 | 0.582494635 | 0.0929 | 0.0851 |
| 1 | 0.6229 | 0.6228 | 0.7892 | 0.586344609 | 0.0922 | 0.0844 |
| 2 | 0.6346 | 0.6475 | 0.8047 | 0.581549654 | 0.0938 | 0.0858 |
| 3 | 0.6365 | 0.6531 | 0.8082 | 0.58779248 | 0.0939 | 0.0855 |
| 4 | 0.6205 | 0.623 | 0.7893 | 0.604152299 | 0.0918 | 0.0834 |
| 5 | 0.6276 | 0.6393 | 0.7995 | 0.587537233 | 0.0927 | 0.0841 |
| 6 | 0.6296 | 0.6441 | 0.8026 | 0.594726828 | 0.0935 | 0.085 |
| 7 | 0.6276 | 0.6415 | 0.8009 | 0.596824932 | 0.0932 | 0.0848 |
| 8 | 0.6493 | 0.676 | 0.8222 | 0.595650904 | 0.0949 | 0.0865 |
| 9 | 0.6235 | 0.6207 | 0.7878 | 0.597243669 | 0.092 | 0.0844 |
| mean | 0.6303 | 0.6405 | 0.8002 | 0.591431724 | 0.0931 | 0.0849 |
| SD | 0.0079 | 0.0159 | 0.0099 | 0.007342073 | 0.0009 | 0.0008 |

**Supplementary Table 3. List of overlapped metabolite-GPCR pairs between interactome network (884 pairs) and experimental assays.**

| Compound | GPCR | UniProt_id | InChi key | Cano_SMILES | GPCR_ML score | mean_value | Activity | Unit | Database |
|---|---|---|---|---|---|---|---|---|---|
| Dexchlorpheniramine | HRH1 | P35367 | SOYKEARSMXGVTM-HNNXBMFYSA-N | CN(C)CCC(C1=CC=C(C=C1)Cl)C2=CC=CC=N2 | 8.02 | 0.73 | Ki | nM | ChEMBL |
| Uridine 5'-Triphosphate | P2Y11 | Q96G91 | PGAVKCOVUIYSFO-XVFCMESISA-N | C1=CN(C(=O)NC1=O)C2C(C(C(O2)COP(=O)(O)OP(=O)(O)OP(=O)(O)O)O)O | 7.95 | 6309.57 | EC50 | nM | IUPHAR |
| Irbesartan | AGTR1 | P30556 | YOSHYTLCDANDAN-UHFFFAOYSA-N | CCCCC1=NC2(CCCC2)C(=O)N1CC3=CC=C(C=C3)C4=CC=CC=C4C5=NNN=N5 | 8.2 | 1.2 | Ki | nM | BindingDB |
| Melatonin | MTR1B | P49286 | DRLFMBDRBRZALE-UHFFFAOYSA-N | CC(=O)NCCC1=CNC2=C1C=C(C=C2)OC | 8.11 | 0.37 | Ki | nM | BindingDB |
